# Contribution of glutamatergic projections to neurons in the nonhuman primate lateral substantia nigra pars reticulata for the reactive inhibition

**DOI:** 10.1101/2024.12.25.630331

**Authors:** Atsushi Yoshida, Okihide Hikosaka

**Author notes:** Correspondence: Atsushi Yoshida. **Author Contributions:** A.Y. contributed to the conception and design of the experiments. A.Y. conducted data collection, analyzed the data, and wrote the manuscript. O.H. supervised this study. **Competing Interest Statement:** The authors declare no competing financial interests.

## Abstract

The basal ganglia play a crucial role in action selection by facilitating desired movements and suppressing unwanted ones. The substantia nigra pars reticulata (SNr), a key output nucleus, facilitates movement through disinhibition of the superior colliculus (SC). However, its role in action suppression, particularly in primates, remains less clear. We investigated whether individual SNr neurons in three male macaque monkeys bidirectionally modulate their activity to both facilitate and suppress actions and examined the role of glutamatergic inputs in suppression. Monkeys performed a sequential choice task, selecting or rejecting visually presented targets. Electrophysiological recordings showed SNr neurons decreased firing rates during target selection and increased firing rates during rejection, demonstrating bidirectional modulation. Pharmacological blockade of glutamatergic inputs to the lateral SNr disrupted saccadic control and impaired suppression of reflexive saccades, providing causal evidence for the role of excitatory input in behavioral inhibition. These findings suggest that glutamatergic projections, most likely from the subthalamic nucleus, drive the increased SNr activity during action suppression. Our results highlight conserved basal ganglia mechanisms across species and offer insights into the neural substrates of action selection and suppression in primates, with implications for understanding disorders such as Parkinson’s disease.

**Significance Statement:** Understanding how the basal ganglia facilitate desired actions while suppressing unwanted ones is fundamental to neuroscience. This study shows that neurons in the primate substantia nigra pars reticulata (SNr) bidirectionally modulate activity to control action, decreasing firing rates to facilitate movements and increasing rates to suppress them. Importantly, we provide causal evidence that glutamatergic inputs to the lateral SNr mediate action suppression. These findings reveal a conserved mechanism of action control in primates and highlight the role of excitatory inputs in behavioral inhibition. This advances our understanding of basal ganglia function and has significant implications for treating movement disorders like Parkinson’s disease.

## Introduction

Our ability to select desired actions while suppressing unwanted ones is essential for daily life, from choosing food to avoiding obstacles. The basal ganglia play a critical role in selective control, forming circuits that regulate action selection and motor control (1–6). A key component of this circuitry is the substantia nigra pars reticulata (SNr), a major output nucleus that tonically inhibits the SC through dense gamma-aminobutyric acid (GABA)-ergic projections (7–11). The SC, vital for controlling saccadic eye movements, is continuously inhibited by the high spontaneous firing rates of SNr neurons (12–19). During saccadic movements, SNr neurons decrease firing, disinhibiting SC neurons and enabling movement execution (20–29). This disinhibition is widely recognized as a fundamental mechanism by which the basal ganglia promote voluntary movements (23, 30–32). While the SNr is known to facilitate desired movements through the disinhibition of downstream targets, its role in suppressing unwanted actions, particularly in primates, remains unclear. Rodent studies suggest increased lateral SNr activity during movement cancellation, possibly driven by excitatory inputs from the subthalamic nucleus (STN) (33–38). However, no study has investigated bidirectional SNr coding in primates during a choice task. This gap raises critical questions: does a similar bidirectional coding scheme operate in primates, with individual SNr neurons facilitating and suppressing actions? What role do excitatory inputs play in suppression within the primate SNr?

In this study, we test the hypothesis that individual SNr neurons in primates bidirectionally modulate activity to facilitate and suppress actions—decreasing during target selection and increasing during rejection. We also propose that excitatory input, particularly glutamatergic projections, predominantly thought to originate from the STN, is critical for mediating suppression within the primate SNr.

To evaluate this, we recorded SNr neuronal activity in macaque monkeys performing a sequential choice task involving selection or rejection of visually presented targets (39). Using electrophysiology and pharmacological blockade of glutamatergic inputs to the lateral SNr, we identified saccade-related neurons. Our findings reveal bidirectional modulation of SNr neurons during the task, with activity decreasing during target selection and increasing during rejection. Blocking glutamatergic input disrupted saccadic control and impaired suppression of reflexive saccades, providing causal evidence for excitatory input’s role in behavioral inhibition.

These findings suggest a conserved basal ganglia function across species and offer new insights into the neural mechanisms of action selection and suppression in primates, with implications for understanding and treating basal ganglia disorders such as Parkinson’s disease.

## Results

### Behavior results in the choice of task

Three monkeys were trained to perform a sequential choice task, evaluating and responding to objects based on learned values (Figure 1A). Monkeys could accept an object by making a saccade and maintaining fixation or reject it through three alternatives: initiating a saccade before returning to the center ("return"), maintaining central gaze without a saccade ("stay"), or making a saccade away ("other") (Figure 1B). Upon rejection, the object disappeared, and a new one was presented until the object was accepted.

**Figure 1.**
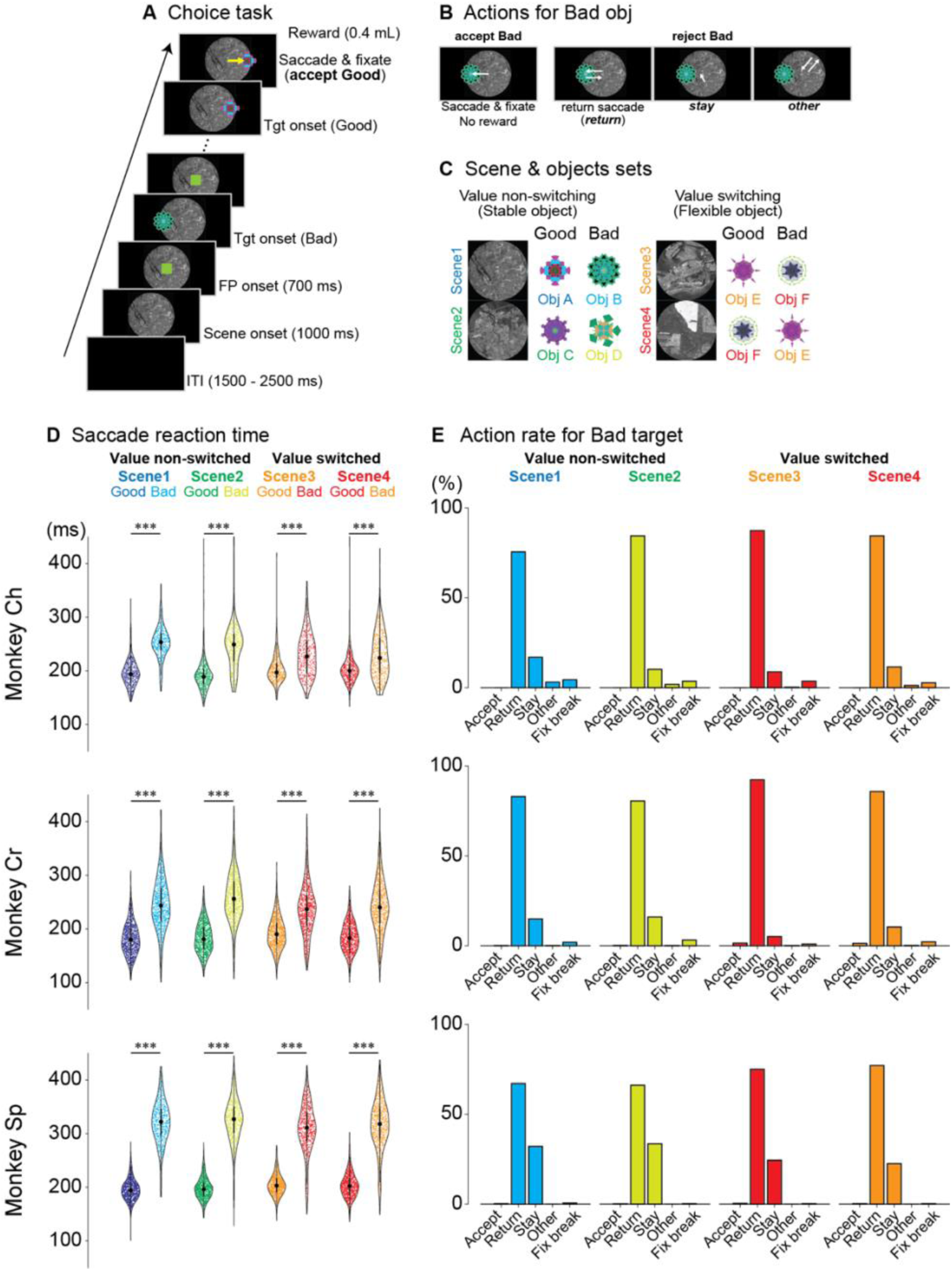
Choice task paradigm and behavioral performance. (A) Time course of the choice task. A background scene is initially presented (1000 ms), followed by a fixation point (FP, 700 ms). Subsequently, either a good (rewarded) or bad (non-rewarded) object is randomly presented at one of six possible locations (0°, 45°, 135°, 180°, 225°, or 315°). (B) Possible behavioral responses to bad objects. When a bad object appears, monkeys can either incorrectly accept it by making a saccade and fixating on it (accept bad, resulting in no reward) or correctly reject it through one of three actions: making a saccade toward the object and immediately returning to center (return), maintaining fixation near the center without making a saccade (stay) or making a saccade in a direction different from the object’s location (other). (C) Scene and object combinations used for neuronal recordings. Each recording session utilized one of six sets, with each set containing four scenes. Each scene was associated with two objects: one good (rewarded) and one bad (non-rewarded). In scenes 1 and 2, object values remained constant (value non-switching), while in scenes 3 and 4, the same objects were used but with reversed reward values (value-switching), illustrating context-dependent value assignment. (D) Saccadic reaction times for three monkeys (Ch, Cr, and Sp). Violin plots show that, across all scenes and monkeys, saccades to good objects had significantly shorter latencies compared to bad objects (***p < 0.0001, Welch’s t-tests). (E) Proportions of different behavioral responses to bad objects across all scenes. All monkeys exhibited similar patterns: incorrect acceptance of bad objects was rare, with rejection responses predominantly consisting of return saccades, followed by stay responses and occasional other responses. Abbreviations: FP, fixation point; ITI, inter-trial interval; Obj, object; Tgt, target.

Each recording session began with one of four randomly selected scenes providing context on the value of two objects: a "good" object yielding a reward and a "bad" object yielding none (Figure 1C). To ensure that neural responses reflected object value and not simply visual features, we implemented a value-switching design in scenes 3 and 4. This involved presenting the same pair of objects (Obj E and Obj F) in both scenes, but reversing their reward contingencies (Obj E good/Obj F bad in scene 3; Obj E bad/Obj F good in scene 4). This allowed us to isolate the effect of value on neural activity while controlling for visual input.

Neural recordings commenced after monkeys achieved stable performance, reliably distinguishing good from bad objects with 90% accuracy across scenes (Figure S1). To avoid bias, one of six object-scene sets was randomly selected for each session (Figure S1).

Saccadic reaction time analysis revealed faster responses to good objects compared to bad ones across all four scenes (Welch’s t-test, *p* < 0.0001; Table S1). This consistent difference indicates that monkeys recognized object values before initiating saccadic responses (Figure 1D).

When rejecting bad objects, monkeys predominantly used the return saccade strategy (Figure 1E), making a saccade toward the bad object and quickly returning their gaze to the center. This accounted for most rejections across all scenes (monkey Ch: 75.6–87.4%; monkey Cr: 80.5– 92.3%; monkey Sp: 66.2–77.1%; see Table S2 for details). The stay strategy, where monkeys maintained central fixation without making a saccade, was less frequent (monkey Ch: 8.7–16.9%; monkey Cr: 5.1–16.0%; monkey Sp: 22.5–33.5%). The preference for the return saccade likely reflects task timing: the stay response required maintaining fixation for 400 ms before the next target appeared, whereas the return response allowed the next target to appear immediately after returning gaze to the center, bypassing the fixed delay. This efficiency likely motivated monkeys to favor the return strategy.

The proportion of stay responses differed between scenes with stable (scenes 1 and 2) versus flexible, context-dependent values (scenes 3 and 4). Monkeys showed significantly more stay responses in scenes 1 and 2 compared to scenes 3 and 4 (Fisher’s exact test: monkey Ch, p < 9.60×10⁻³, φ = 0.06, 95% CI: 0.53–0.92; monkey Cr, p < 9.68×10⁻¹⁴, φ = 0.12, 95% CI: 0.38–0.58; monkey Sp, p < 5.22×10⁻¹⁰, φ = 0.11, 95% CI: 0.54–0.73). This difference may reflect the higher cognitive demand of processing context-dependent values in scenes 3 and 4.

### Neuronal activity of SNr neurons during the choice task

To investigate how SNr neurons encode object evaluation and action selection, we recorded single-unit activity from 129 neurons across three monkeys (29 from monkey Ch, 51 from monkey Cr, and 49 from monkey Sp) during the choice task. Neurons were included based on observed activity changes following target onset, specifically those with significant firing rate modulation.

A representative SNr neuron showed distinct activity patterns in response to good and bad objects (Figures 2A and 2B). While minimally responsive to scene onset (Figure 2A), this neuron exhibited robust value-dependent modulation after target presentation: firing rates decreased for good objects and increased for bad objects across all scenes (Figure 2B). This bidirectional response pattern was consistent for both contralateral and ipsilateral target presentations.

**Figure 2.**
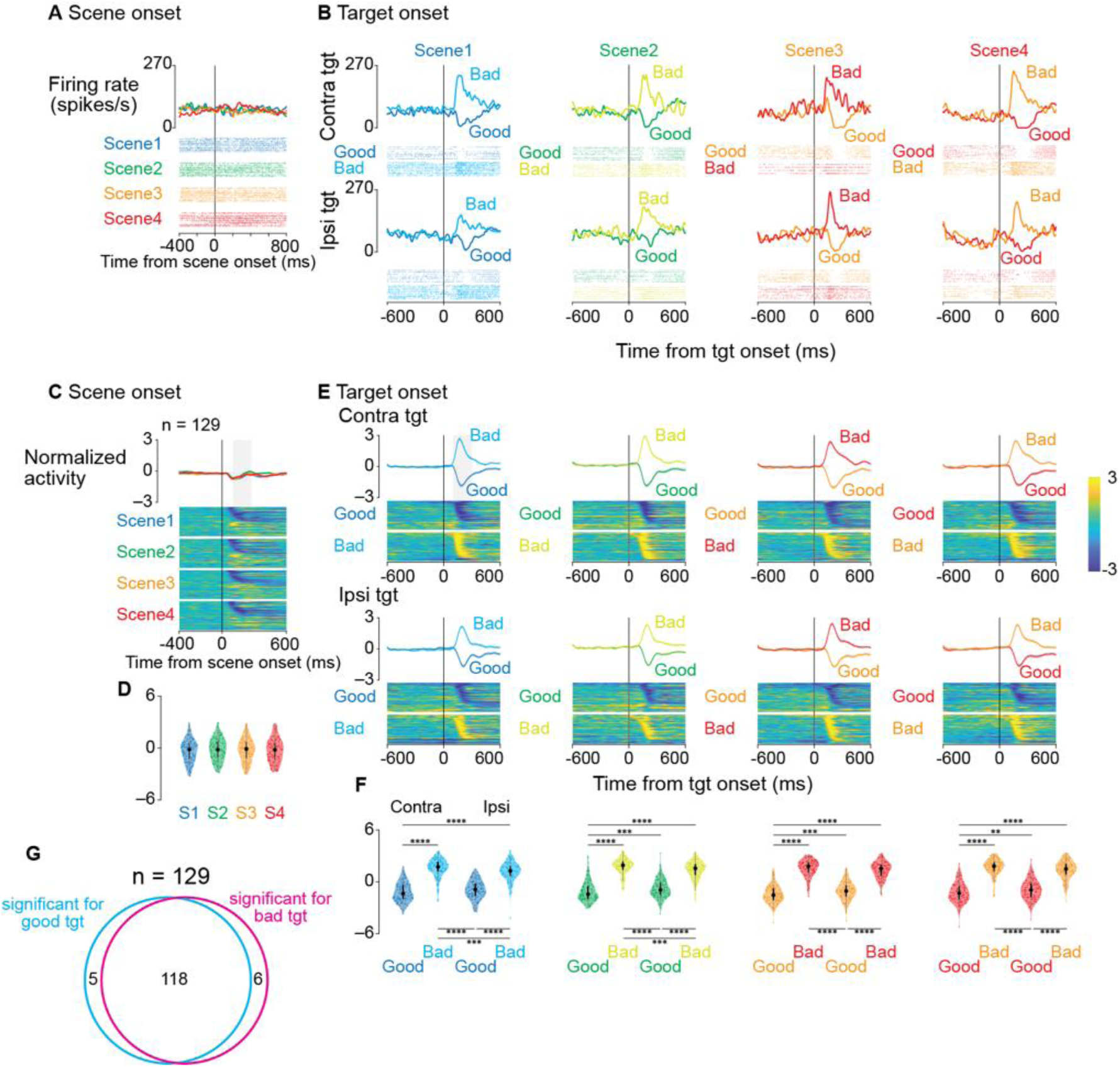
Single-neuron and population responses to scene and target presentation during the choice task. (A, B) Activity of a representative SNr neuron. Spike density functions (top) and raster plots (bottom) are shown aligned to scene onset (A) and target onset (B). In (A), different colors represent different scenes. In (B), activity is displayed separately for contralateral (upper) and ipsilateral (lower) target presentations, with different colors indicating responses to good and bad objects in each scene. (C) Population activity aligned to scene onset (n = 129 neurons). The upper panels show mean firing rates (±SEM, shaded area) for each scene. Lower panels display color-coded normalized firing rates of individual neurons (rows) over time. (D) Distribution of neuronal responses to scene onset. Violin plots quantify activity during a 200-ms window starting 100 ms after scene onset (gray rectangle in [C]). Plot elements represent the median (large circle), interquartile range (thick line), and range (thin line). (E) Population activity aligned to target onset, shown separately for contralateral and ipsilateral presentations of good and bad objects across all scenes. The format is consistent with (C). (F) Distribution of neuronal responses to target presentation. Violin plots quantify activity during a 200-ms window starting 100 ms after target onset. Asterisks denote significant differences between conditions (*p < 0.05, **p < 0.01, ***p < 0.001, ****p < 0.0001, post-hoc pairwise t-tests with Bonferroni correction). (G) Venn diagram depicting the distribution of neurons with significant response modulation to target presentation in scene 1 (contralateral targets). Neural activity was compared between a 200-ms pretarget baseline period and a 200-ms window starting 100 ms after target onset. Of 129 neurons, 118 showed significant modulation to both good and bad objects, while 5 and 6 neurons responded exclusively to good or bad objects, respectively. Abbreviation: SEM, standard error of the mean.

Population analyses confirmed this response pattern as characteristic of SNr neurons (Figures 2C and 2E). Normalized population activity showed minimal modulation at scene onset (Figure 2C) but exhibited strong value-dependent responses following target presentation (Figure 2E): decreased activity for good objects and increased activity for bad objects. Analysis within a 200-ms window (100-300 ms after target onset) revealed significant differences between good and bad object responses (parametric bootstrap tests for linear mixed-effects models with post-hoc pairwise t-tests, Bonferroni-corrected; see Table S3). Neural responses did not differ significantly across scenes (parametric bootstrap tests, full vs. null models; p = 0.44), suggesting activity reflected object values rather than visual features. This conclusion is supported by scenes 3 and 4, where identical objects elicited opposite responses based on context-dependent values.

Most SNr neurons exhibited bidirectional value coding. In scene 1, 91.5% (118/129) of neurons showed significant modulation to both good and bad objects during a 200-ms window (100-300 ms after target onset) compared to a 200-ms pretarget baseline (two-sample t-test), while only 3.9% (5/129) and 4.7% (6/129) responded exclusively to good or bad objects, respectively (Figure 2G). This suggests that individual SNr neurons dynamically encode both facilitation and suppression of actions through bidirectional modulation.

To assess whether SNr activity after target onset was related to saccade initiation, we realigned responses to saccade onset (Figure S2). The neuron in Figure 2A exhibited distinct temporal patterns for good versus bad objects (Figure S2A). For good objects, firing rates reached a minimum near saccade initiation, while for bad objects, peak activity occurred before saccade onset.

This pattern was consistent at the population level (Figure S2B). When accepting good objects, population activity showed maximal suppression around saccade initiation. In contrast, for bad objects (return saccades), peak activation occurred before movement onset. Statistical analysis confirmed significant differences between these peri-saccadic response patterns (parametric bootstrap tests for linear mixed-effects models with Bonferroni-corrected post-hoc t-tests; see Table S4).

To examine the link between SNr activity and saccadic reaction times, we analyzed correlations between normalized firing rates (150 ms before to 50 ms after saccade onset) and normalized reaction times (Figure S2D). Reaction times were normalized due to skewed distributions for Bayesian linear mixed-effects models. A significant positive correlation was found only for contralateral bad objects (posterior mean β = 0.075, 95% credible interval [0.01, 0.14], P(β > 0) = 99.2%), while no significant correlations were observed for other conditions (contralateral good, ipsilateral good, ipsilateral bad; see Table S5). The correlation between increased SNr activity and longer reaction times for contralateral bad objects aligns with the inhibitory role of SNr projections to the SC: stronger SNr activation likely enhances SC suppression, delaying saccade initiation for contralateral targets.

### Neuronal activity in the SNr during choice rejection and saccade suppression

To investigate the role of SNr in behavioral inhibition, we compared neuronal responses across rejection strategies and behavioral contexts. Monkeys rejected bad objects by either making a saccade toward the object and returning to the center ("return") or maintaining central fixation without a saccade ("stay") (Figure 3A). While both strategies resulted in rejection, "return" involved saccade execution, and "stay" required saccade suppression.

**Figure 3.**
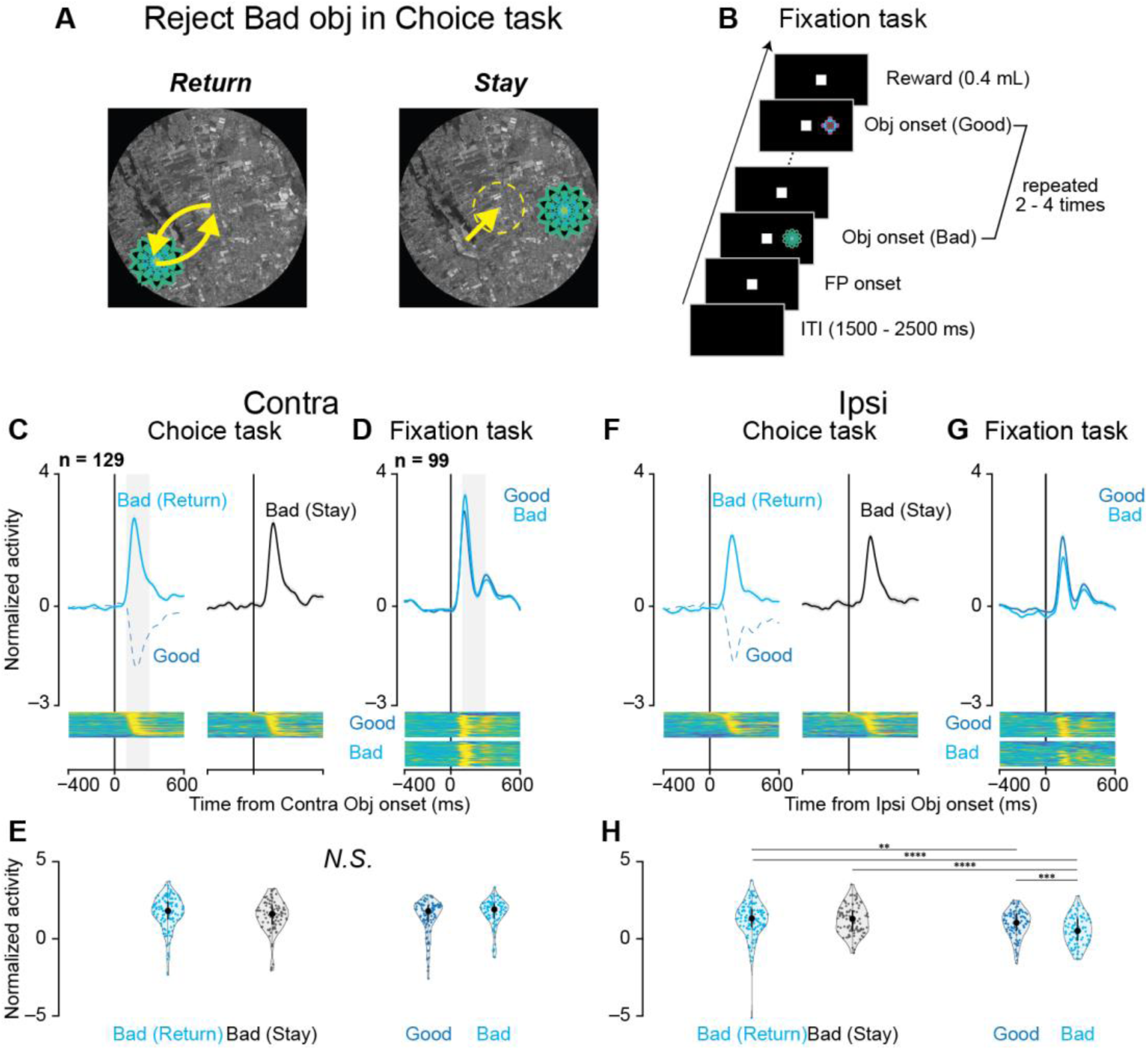
Neural activity during different rejection strategies and saccade suppression. (A) Schematic illustration of two rejection strategies for bad objects: "return" (saccade toward the object followed by return to center) and "stay" (maintaining central fixation). (B) Fixation task design. Objects from scene 1 of the choice task (both good and bad) were presented sequentially (2–4 presentations) in the contralateral or ipsilateral visual field. Monkeys were rewarded for maintaining central fixation throughout the presentations, requiring active suppression of reflexive saccades to the objects. (C, F) Population responses during bad object rejection in scene 1 are shown separately for contralateral (C) and ipsilateral (F) target presentations. Cyan traces represent "return" responses, black traces represent "stay" responses, and blue dotted traces represent responses during good object acceptance for comparison. Lower panels display color-coded normalized firing rates of individual neurons. (D, G) Population responses during the fixation task for contralateral (D) and ipsilateral (G) object presentations. Blue traces represent responses to good objects, and cyan traces represent responses to bad objects. Lower panels show normalized single-neuron responses as in (C, F). (E, H) Distribution of neuronal responses during the choice task (E) and fixation task (H). Violin plots quantify activity during a 200-ms window starting 100 ms after object onset (gray rectangles in C, D). For ipsilateral presentations (H), asterisks indicate significant differences between conditions (*p < 0.05, **p < 0.01, ***p < 0.001, ****p < 0.0001, post-hoc pairwise t-tests with Bonferroni correction). No significant differences were observed for contralateral presentations (E).

Population activity during bad object rejection showed similar patterns between "return" and "stay" responses for both contralateral (Figure 3C) and ipsilateral (Figure 3F) presentations. Quantitative analysis of neuronal responses (200-ms window, 100–300 ms after object onset) revealed no significant differences between strategies (parametric bootstrap tests for linear mixed-effects models with Bonferroni -corrected post-hoc t-tests; Figures 3E and 3H).

To examine SNr involvement in saccade suppression, we recorded 99 of the 129 neurons during a fixation task (Figure 3B). Monkeys suppressed reflexive saccades to sequentially presented objects (2–4 presentations) while maintaining central fixation to earn a reward. The same objects from scene 1 of the choice task were used in the fixation task to control for visual features and isolate the effect of behavioral context.

During the fixation task, SNr neurons exhibited increased activity following object presentation in both contralateral (Figure 3D) and ipsilateral (Figure 3G) directions. Notably, this increase occurred even for objects labeled as "good" in the choice task, which had previously reduced SNr activity and facilitated saccadic responses (dotted lines in Figures 3C and 3F). The activity increase during central fixation in the fixation task was comparable to that observed during rejection in the choice task, with no significant differences for contralateral presentations (Figure 3E) and some significant differences for ipsilateral comparisons (Figure 3H; see Table S6 for details).

Comparing the fixation and choice tasks highlights their distinct behavioral demands. The fixation task primarily challenges the monkey to reactively inhibit reflexive saccades to suddenly presented objects without evaluating their value. In contrast, the choice task involves a more complex process: reactive inhibition allows time to assess object value, followed by proactive inhibition (rejecting undesirable objects) or rapid disinhibition (accepting desirable objects). This comparison underscores that the fixation task isolates basic reactive suppression, while the choice task builds on this mechanism to incorporate value-based decision-making and guide action selection.

### Effects of glutamatergic receptor antagonists on the SNr

Our neurophysiological findings suggest that increased SNr activity during both proactive inhibition (rejecting bad objects in the choice task) and reactive inhibition (suppressing reflexive saccades in the fixation task) may be driven by excitatory inputs to the SNr. To test this hypothesis, we performed local pharmacological manipulation of glutamatergic transmission in the lateral SNr, where most task-related neurons were located. We injected glutamate receptor antagonists (a mixture of N-methyl-d-aspartate (NMDA) receptor antagonist (carboxypiperazin-4-propyl-1-phosphonic acid, CPP) and aminomethylphosphonic acid (AMPA) receptor antagonist (2,3-dihydroxy-6-nitro-7-sulfamoyl-benzo (F)quinoxaline, NBQX)) into the lateral SNr while monkeys performed both the choice and fixation tasks. The effects of this manipulation were striking and direction-specific. In the choice task, reaction times for contralateral saccades decreased significantly for both good and bad objects (Figure 4A; *p* < 0.0001 for both comparisons, parametric bootstrap tests for generalized linear mixed-effects models with Bonferroni correction, see Table S7). More critically, the manipulation altered rejection patterns: for contralateral bad objects, monkeys showed more "return" responses and fewer "stay" responses, while the opposite pattern occurred for ipsilateral bad objects (Figure 4B; *p* < 0.0001 for all comparisons, see Table S8).

**Figure 4.**
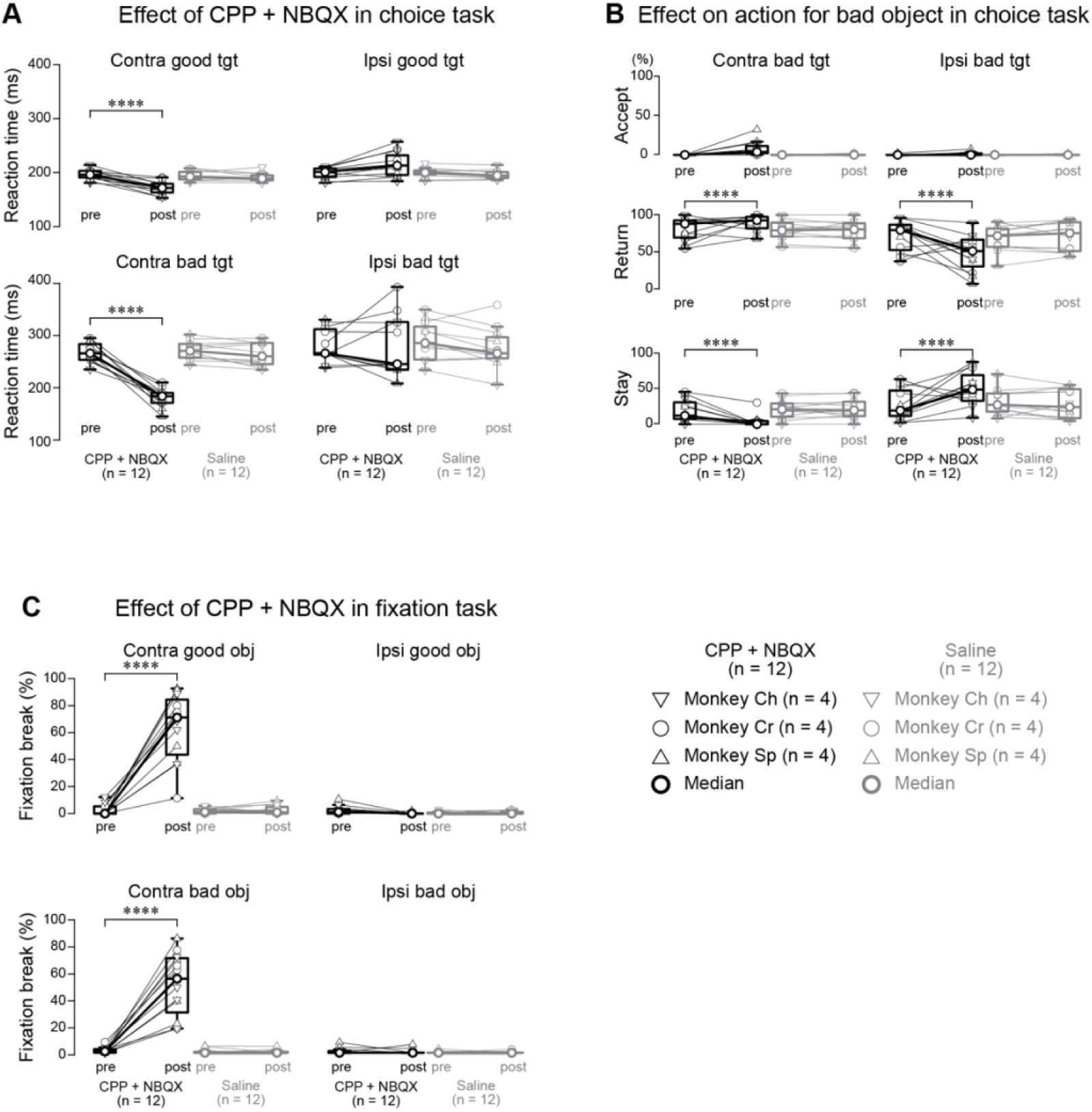
Behavioral effects of glutamatergic receptor antagonists during choice and fixation tasks. (A) Saccadic reaction times before (pre) and after (post) local injection of CPP/NBQX mixture or saline. Data points represent median values for individual sessions from three monkeys (inverted triangles: monkey Ch; circles: monkey Cr; triangles: monkey Sh; n = 4 sessions per condition per monkey). Thick circles and lines indicate population medians. Following CPP/NBQX injection, reaction times significantly decreased for both good and bad objects presented contralaterally. (B) Proportions of different actions (accept, return, stay) toward bad objects before and after drug injection. Format as in (A). CPP/NBQX injection significantly increased the proportion of return responses while decreasing stay responses for contralateral targets, with the opposite pattern observed for ipsilateral targets. (C) Fixation break error rates during the fixation task before and after injection. Format as in (A). CPP/NBQX significantly increased fixation break errors for both good and bad objects presented contralaterally. Asterisks denote significant differences (*p < 0.05, **p < 0.01, ***p < 0.001, ****p < 0.0001, post-hoc pairwise t-tests with Bonferroni correction). Abbreviations: CPP, (±)-3-(2-carboxypiperazin-4-yl) propyl-1-phosphonic acid (NMDA receptor antagonist); NBQX, 2,3-dihydroxy-6-nitro-7-sulfamoyl-benzo[f]quinoxaline (AMPA receptor antagonist); Tgt, target.

The most prominent effect occurred in the fixation task, where glutamate receptor blockade severely impaired reflexive saccade suppression. After antagonist injection, monkeys showed a significant increase in fixation break errors for contralateral object presentations (Figure 4C; *p* < 0.0001, see Table S9). These effects were specific to glutamate receptor blockade, as saline injections caused no significant behavioral changes in either task (Tables S7–S9).

Blocking glutamatergic transmission in the lateral SNr affected saccadic reaction times and altered rejection behaviors, particularly for contralateral targets. The lateralized effects in both tasks underscore the SNr’s role in saccadic control.

These findings suggest a two-stage inhibition process. Reactive inhibition in the lateral SNr initially suppresses reflexive saccades to new objects, providing time to evaluate the object’s value. After value assessment, proactive inhibition (for bad objects) or disinhibition (for good objects) determines the monkey’s next action—accepting good objects quickly or rejecting bad ones via return or stay strategies. Reactive suppression thus facilitates value-based decision-making, ensuring saccadic responses are not reflexive but guided by learned associations.

### Visualization of neuronal recording and injection sites using quantitative susceptibility mapping (QSM)

To localize our recording and injection sites within the SNr, which is challenging to visualize with conventional T1- and T2-weighted magnetic resonance imaging (MRI) due to its signal similarity to surrounding structures, we used QSM, an advanced MRI technique that exploits the paramagnetic properties of iron-rich regions (Figure 5A). As shown in prior work (40), QSM provides clear visualization of subcortical structures, including the iron-rich SNr, in macaques. Our QSM pipeline included multi-echo gradient echo acquisition, phase unwrapping, background field removal, and dipole field inversion to create susceptibility maps.

**Figure 5.**
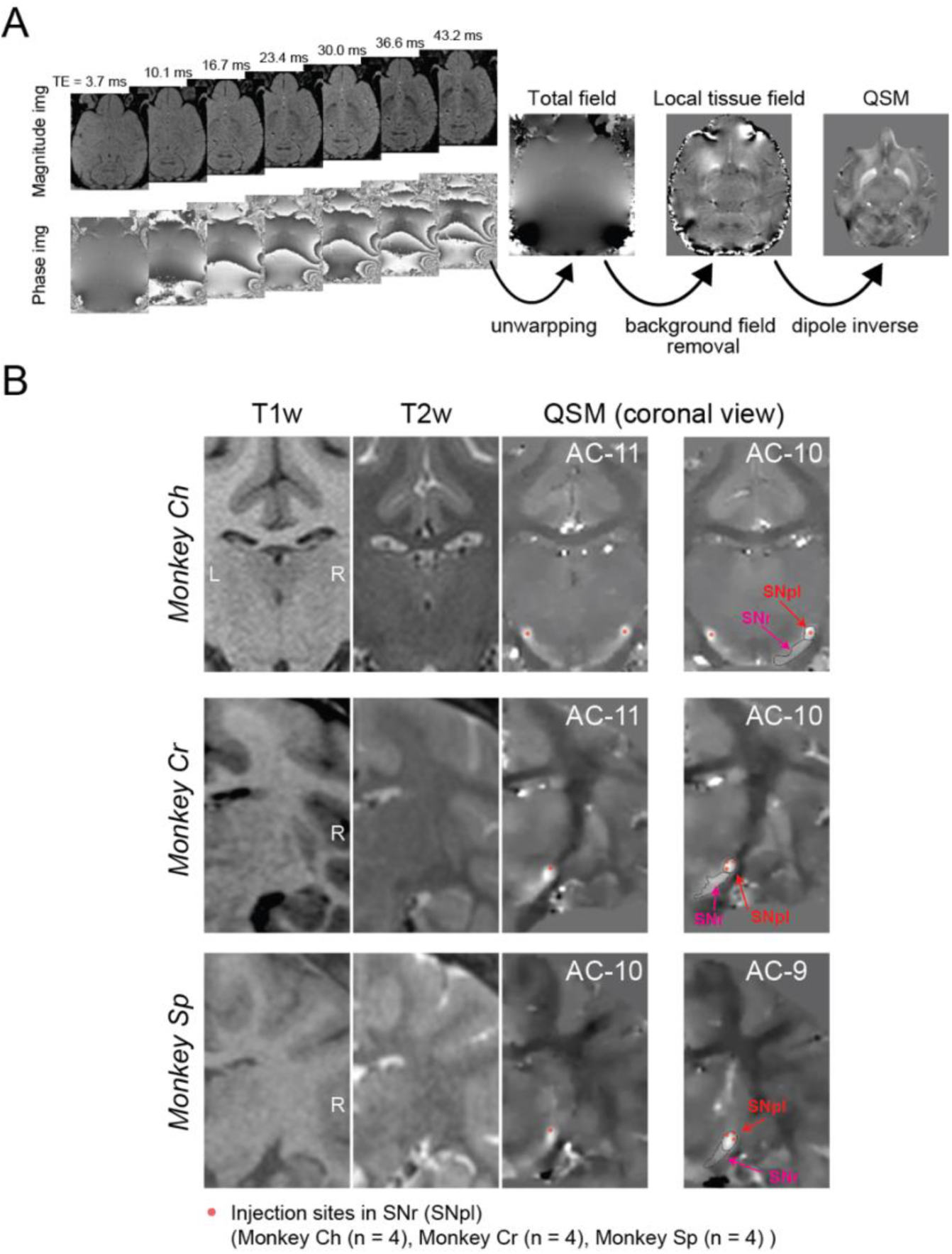
Visualization of injection sites using quantitative susceptibility mapping (QSM) (A) QSM image processing pipeline. Multi-echo 3D gradient echo sequences were acquired with seven echo times (TE: 3.7–43.2 ms). The figure displays magnitude and phase images at different TEs, along with the total field map, local tissue field map, and final QSM image. The processing stream involves three main steps: phase unwrapping of the raw phase images, background field removal using high-pass filtering, and dipole deconvolution to generate the final QSM images. (B) Anatomical localization of injection sites in the substantia nigra pars reticulata (SNr) for all three monkeys. Coronal sections show T1-weighted, T2-weighted, and QSM images at different anterior-posterior levels (relative to the anterior commissure, AC). Red dots mark injection sites within the SNr (lateral part, SNpl). Images are provided for monkey Ch (AC-11 to AC-10), monkey Cr (AC-11 to AC-10), and monkey Sp (AC-10 to AC-9), with n = 4 injection sites per monkey. Abbreviations: AC, anterior commissure; QSM, quantitative susceptibility mapping; SNpl, substantia nigra pars lateralis; SNr, substantia nigra pars reticulata; TE, echo time.

QSM offered superior visualization of basal ganglia structures compared to T1- and T2-weighted imaging (Figure 5B). High susceptibility values were observed in the lateral SNr, corresponding to the substantia nigra pars lateralis (SNpl), which showed the strongest QSM signal and the highest concentration of task-related neurons. This region also served as the site for targeted pharmacological manipulations. The enhanced contrast in QSM images allowed precise confirmation of recording and injection sites, ensuring the anatomical specificity of our experimental interventions.

## Discussion

We examined the role of the substantia nigra pars reticulata (SNr) in action selection and suppression, focusing on glutamatergic inputs. We addressed two major questions: (1) whether individual SNr neurons in primates bidirectionally modulate activity to facilitate and suppress actions, and (2) whether glutamatergic input provides the causal drive for these inhibitory processes. Using electrophysiological recordings, behavioral tasks, and pharmacological manipulations, we provided clear answers to these questions. Recordings from SNr neurons in macaque monkeys during a sequential choice task revealed bidirectional coding: neurons decreased firing rates for desirable targets and increased firing for undesirable ones. This dynamic modulation was consistent across contexts and linked to monkeys’ choices, highlighting the SNr’s role in both facilitating and suppressing actions. Pharmacological blockade of glutamatergic inputs to the lateral SNr disrupted saccadic control, altering reaction times and rejection behaviors. These findings demonstrate a causal role for excitatory inputs in behavioral inhibition and address key gaps in understanding SNr function.

### Does the lateral SNr mediate reactive, proactive, or both forms of inhibition?

Behavioral inhibition is classified into proactive and reactive forms (41). Proactive inhibition involves anticipatory action suppression to achieve a goal, enabling planned restraint. Reactive inhibition is the immediate suppression of actions in response to unexpected stimuli, allowing rapid adjustment to environmental changes. In our sequential choice task, rejecting a bad object represents proactive inhibition, as it serves the goal of accepting a good object for a reward. However, our findings indicate that the lateral SNr primarily supports reactive, not proactive, inhibition.

Glutamate receptor antagonist injections into the lateral SNr significantly shortened saccadic reaction times to *both* contralateral good *and* bad objects in the choice task (Figure 4A). If the lateral SNr were solely responsible for proactive inhibition, antagonist injections would likely impair bad object rejection, leading to *more* saccades toward or acceptance of bad objects. Instead, we observed faster saccades to *both* good and bad objects, suggesting a disruption of reactive inhibition, which normally suppresses reflexive saccades to sudden stimuli. Blocking glutamatergic input to the SNr likely weakened this reactive suppression, resulting in faster but less controlled saccades.

We propose that reactive and proactive inhibition both operate in the choice task but serve distinct roles. Reactive inhibition initially suppresses reflexive saccades to all presented objects, providing a brief window for object evaluation before a response is made. If the object is deemed bad, proactive inhibition engages to reject it.

This hypothesis is supported by saccadic reaction times (Figure 1D). Across all monkeys, saccades to good objects were faster than to bad objects, indicating that object evaluation occurs *before* saccade initiation. If evaluation followed reflexive saccades, no reaction time difference would be expected.

The timing of lateral SNr activity further supports this model. During bad object rejections, peak SNr activity *precedes* return saccade onset (Figure S2), suggesting a role in anticipating movement suppression (reactive inhibition) rather than executing proactive inhibition. This contrasts with findings in the anterior striatum in our previous study (39), where proactive inhibition-related activity decreases *after* return saccade initiation, emphasizing distinct roles for the SNr in reactive and the striatum in proactive inhibition.

The fixation task and comparisons between "return" and "stay" responses reinforce the focus on reactive inhibition. In the fixation task, glutamate antagonist injections increased fixation break errors, highlighting the importance of glutamatergic inputs to the SNr for reactive suppression. During the choice task, no significant difference in SNr activity was observed between "return" and "stay" responses to bad objects (Figure 3), indicating that increased activity reflects reactive suppression of reflexive saccades rather than specific motor execution. Similar increases in SNr activity during fixation and choice tasks, particularly for contralateral presentations, strongly support the lateral SNr’s primary role in reactive inhibition.

### Potential sources of excitatory input to the lateral SNr

Identifying the source of excitatory inputs to the lateral SNr that mediate behavioral inhibition is crucial for interpreting our findings. Blocking glutamatergic transmission in the SNr shortened saccadic reaction times in the choice task and increased fixation break errors in the fixation task, emphasizing the importance of excitatory projections to the lateral SNr.

The subthalamic nucleus (STN) is the most likely source of these inputs. Strong anatomical and physiological evidence supports a direct STN-SNr projection, a critical component of both the indirect and hyper-direct basal ganglia pathways (42,43). Optogenetic studies in rodents also implicate STN excitatory projections in behavioral inhibition (38).

Other sources cannot be definitively excluded. Evidence suggests potential direct projections from the motor cortex to the SNr in rodents (44), raising the possibility that cortical inputs might also contribute to the SNr’s excitatory drive. However, the existence and significance of such a cortico-SNr pathway in primates remain unclear. Future studies using selective optogenetic manipulation or pathway-specific tracing could clarify the relative contributions of the STN and motor cortex to the lateral SNr’s excitatory inputs in primates.

### Comparative analysis of lateral SNr function: Conserved bidirectional coding?

Our study revealed that most lateral SNr neurons (91.5%, 118/129) exhibit bidirectional coding, modulating activity to encode both saccade facilitation and suppression. This aligns with previous findings in other species. For instance, Schmidt et al. observed that 55.6% (10/18) of rat lateral SNr neurons decreased activity during planned actions and increased activity during successful cancellations in a stop-signal task (33, 45). This suggests that bidirectional modulation related to action control may be evolutionarily conserved.

In contrast, Jiang et al., studying anesthetized cats, reported distinct SNr neuron populations projecting to the ipsilateral and contralateral SC, with inhibitory and excitatory responses to contralateral visual stimuli, respectively (46). This suggests functional segregation of SNr neurons rather than bidirectional coding within individual neurons. However, anesthesia profoundly alters neuronal activity and synaptic transmission, potentially masking true functional properties. Differences in experimental conditions (anesthetized cats vs. awake rats and monkeys in our study) likely account for these discrepancies.

These contrasting results emphasize the need to consider species differences and experimental conditions when studying SNr function. The disparity between findings in awake and anesthetized animals highlights how consciousness can profoundly influence neural circuit dynamics. Further research in awake preparations across multiple species is essential to determine whether bidirectional coding in the lateral SNr is a conserved feature and to clarify the effects of anesthesia on SNr responses. This will enhance our understanding of the principles governing SNr function in action control. The evolutionary origins and conservation of bidirectional coding in the SNr remain open questions. Investigations using advanced techniques like optogenetics or high-resolution imaging in primate models are crucial to unravel these mechanisms.

## Materials and Methods

The behavioral tasks and experimental procedures were identical to those in our previous investigation of striatal activity during choice and fixation tasks (39). Below, we briefly outline essential procedures and refer readers to our earlier work for additional details.

### Animal preparation

All procedures were approved by the National Eye Institute Animal Care and Use Committee and complied with the Public Health Service Policy on Laboratory Animal Care. Three male macaque monkeys (Macaca mulatta, 8–10 kg), referred to as Monkeys Ch, Cr, and Sp, were used. Monkeys Cr and Sp also participated in our previous study (39). Under isoflurane anesthesia and sterile conditions, we implanted a plastic head holder and recording chambers. After recovery, monkeys were trained on oculomotor tasks. During experimental sessions, their heads were fixed, and eye movements were tracked using an infrared device (EyeLink 1000, SR Research) at 1,000 Hz. Water intake was regulated to maintain task motivation. For detailed surgical and postoperative procedures, see our previous report (39).

### Behavioral results

Experiments were conducted in a dark, soundproof room. Visual stimuli were presented on a screen using an LCD projector (PJ658, ViewSonic). Task control and data acquisition were managed using custom software (Blip; http://www.robili.sblip/). As in our previous study (39), we used two behavioral tasks: a choice task and a fixation task.

### Choice task

The structure and timing of the choice task matched our previous study. Briefly, one of six scene-object sets was randomly selected for each recording session (Figure S1). Each set included four scenes with two fractal objects per scene: one "good" (rewarded) and one "bad" (non-rewarded). In scenes 1 and 2, object values were stable, while in scenes 3 and 4, values were reversed, dissociating visual features from value coding.

On each trial, after scene presentation (1,000 ms) and central fixation (700 ms), either a good or bad object appeared at one of six peripheral locations (15° eccentricity). Monkeys could accept the object by making a saccade and maintaining fixation (>400 ms) or reject it through three behaviors: returning to center after a brief saccade ("return"), maintaining central fixation ("stay"), or looking away ("other"). Accepting good objects earned juice rewards (0.4 mL). After rejection, another object was presented until acceptance.

### Fixation task

After isolating task-related neurons in the choice task, we conducted a fixation task to examine SNr activity in saccade suppression. Good and bad objects from scene 1 of the choice task were presented sequentially (2–4 times, 400 ms duration, 400 ms interval) while monkeys maintained central fixation. Successful fixation through the sequence earned a reward, while saccades toward objects were counted as fixation break errors.

### MRI

After implanting recording chambers, an MRI was performed to map cerebral structures and grid apertures. A gadolinium-enhanced contrast medium (Magnevist, Bayer Healthcare Pharmaceuticals) was introduced into the chambers. Recording sites were identified using a high-definition 3T MR system (MAGNETOM Prisma; Siemens Healthcare) with three-dimensional T1-weighted (T1w, MPRAGE) and T2-weighted (T2w, SPACE) sequences (0.5 mm voxel size).

Because conventional T1w and T2w images provided poor contrast between the SNr and surrounding structures, QSM was used to enhance SNr visualization (47, 48, 40). QSM images were reconstructed from phase images acquired with a 3D multi-echo gradient echo sequence (repetition time: 50 ms; echo times: 3.7, 10.1, 16.7, 23.4, 30.0, 36.6, 43.2 ms) (40). The QSM reconstruction pipeline included multiple steps:

1. Phase Unwrapping: Phase images were unwrapped.
2. Background Field Removal: High-pass filtering was applied to the phase images.
3. Dipole Inversion: Dipole field inversion was performed to generate quantitative susceptibility maps.

These steps were implemented using the morphology-enabled dipole inversion toolbox (http://pre.weill.cornell.edu/mri/pages/qsm.html) (49) in MATLAB 2019 (The MathWorks, Inc., Natick, MA, USA).

### Neuronal recording procedure

Single-unit activity was recorded from the SNr using tungsten electrodes (1–9 MΩ; Frederick Haer & Co.; Alpha Omega Engineering). Electrodes were advanced through stainless steel guide tubes with a hydraulic micromanipulator (MO-973A, Narishige). Neural signals were amplified, bandpass filtered (0.3–10 kHz; A-M Systems), and digitized at 40 kHz. Individual neurons were isolated online using custom voltage-time window discrimination software (Blip). Recording began once stable isolation was achieved, and the neuron showed activity modulation following target onset in the choice task.

### The glutamatergic antagonist injection procedure

To inhibit excitatory projections to the SNr, we injected a mixture of CPP (C104, Sigma-Aldrich) and NBQX (N183, Sigma-Aldrich) into the lateral SNr, where task-related neurons were concentrated. The high baseline firing rate of SNr neurons allowed the precise localization of the dorsal SNr edge with the electrode tip. Prior to injection, neuronal activity was recorded using a custom injectrode (50).

Monkeys first performed the choice and fixation tasks to collect preinjection control data. Then, 1 μL of a 5–10 mM mixture of CPP and NBQX was injected at 0.2 μL/min using a remotely controlled infusion pump (PHD ULTRA, Harvard Apparatus). Concentrations were based on a previous study (51), with higher levels used to assess the effects of excitatory projections on behavior. After injection, monkeys repeated the choice and fixation tasks to evaluate the effects of CPP and NBQX (5–90 min postinjection). As a control, 1 μL of saline was injected into the same SNr sites.

For statistical analyses, data from 60–90 min postinjection of CPP, NBQX, and saline were used. However, for one data point from monkey Sp (10 mM CPP and NBQX), analysis was performed at 20 min postinjection because the effects were too strong for task continuation.

### Data analysis and Statistical analysis

All behavioral and neurophysiological data were preprocessed using MATLAB 2022b (MathWorks, Natick, MA, USA). The sample size was not predetermined statistically but guided by previous studies on recorded SNr neurons (28).

### Behavior data analysis

In the choice task, saccade onsets were defined as eye velocity exceeding 40°/s within 400 ms of target onset. Reaction times for good and bad objects (Figure 1D) included only initial saccades toward objects, excluding return movements. Welch’s t-tests compared reaction times across conditions. Fisher’s exact test was used to compare "stay" responses between stable (scenes 1 and 2) and flexible (scenes 3 and 4) value conditions (Figure 1E). In the fixation task, saccades toward presented objects were classified as fixation break errors.

### Neuronal data processing

Neuronal data were aligned with event initiation (scene, target, and saccade onset). Peristimulus time histograms (PSTHs) were calculated in 1-ms bins and smoothed with a Gaussian filter (σ = 20 ms). Neuronal activity was Z-transformed by subtracting the baseline firing rate (average firing during the 500ms before event onset) from the smoothed PSTH and dividing it by the PSTH’s SD. The time course of responses was analyzed for each condition using these Z-transformed PSTHs.

To quantify neurons showing significant modulation to good and/or bad objects (Figure 2G), neural activity was compared between baseline and post-stimulus periods for contralateral targets in scene 1. The baseline period was 200ms before the target onset, and the post-stimulus period was 200ms starting 100 ms after the target onset. For each neuron, two-sample t-tests (α = 0.05) compared normalized firing rates between these periods for good and bad objects. Neurons were classified as responding to: (1) good objects only, (2) bad objects only, or (3) both. These classifications were visualized using a Venn diagram (Figure 2G).

### Statistical modeling

To compare normalized neuronal activity and behavioral parameters (e.g., saccade reaction times) before and after injections, we used linear mixed-effects models (LMMs) and generalized linear mixed-effects models (GLMMs). These hierarchical models incorporate random effects for individual subjects and neurons, accounting for repeated measurements. By modeling fixed and random effects, LMMs/GLMMs reduce type I errors and better represent the data structure (54). LMMs were used to analyze Z-transformed SNr neuronal activity under various conditions (Figures 2, 3, and S2). GLMMs compared saccade reaction times, proportions of chosen actions for bad objects in the choice task, and fixation break errors in the fixation task (Figure 4) before and after injection. For all LMM and GLMM tests, we compared full models with explanatory variables as fixed effects and random effects for monkey, neuron, or session IDs (null model). Detailed model specifications are provided in Supplementary Methods.

We used a parametric bootstrap method to assess model goodness of fit, performing 10,000 iterations and computing the p-value from the deviance difference between models. If the full model demonstrated significant fit, post-hoc pairwise t-tests with Bonferroni correction were conducted to explore differences. For LMM and GLMM analyses, we used the lme4 (55), pbkrtest (56), emmeans (57), and brms (58) packages in RStudio.

## Supplementary Methods

### Detailed model specifications

Model specifications for the statistical analyses presented in the main text are detailed below. All linear mixed-effects (LMM) and generalized linear mixed-effects (GLMM) models included random effects (e.g., [1|monkey_ID] or [1|monkey_ID: Neuron_ID]) to statistically control for individual differences at both the subject and neuron levels. By accounting for these hierarchical structures, the models minimized the risk of inflated type I errors due to the nonindependence of observations within the same monkey or neuron, ensuring more accurate and robust inferences. All models were fit using RStudio with the following packages: lme4 (Bates et al., 2014), pbkrtest (Halekoh & Højsgaard, 2014), emmeans (Lenth et al., 2019), and brms (Bürkner, 2017).

### 1. Linear mixed-effects model for neuronal activity at scene onset (Figure 2D)

Model objective: To evaluate differences in neuronal activity following scene onset.

Model specifications:

Full Model: NormalizedNeuronalActivity ∼ Scene + (1|monkey_ID) + (1|monkey_ID: Neuron_ID) Null Model: NormalizedNeuronalActivity ∼ (1|monkey_ID) + (1|monkey_ID: Neuron_ID)

Variables:

NormalizedNeuronalActivity: Mean Z-transformed PSTH (100-300 ms after scene onset) Scene: Fixed effect (levels: 1-4)

monkey_ID, Neuron_ID: Random effects

Statistical threshold: α = 0.05

### 2. Linear mixed-effects model for neuronal activity at target onset (Figure 2F)

Model objective: To examine how neuronal activity varies with scene context, object value, and target direction.

Model specifications:

Full Model: NormalizedNeuronalActivity ∼ Scene × Value × Direction + (1|monkey_ID) + (1|monkey_ID: Neuron_ID)

Null Model: NormalizedNeuronalActivity ∼ (1|monkey_ID) + (1|monkey_ID: Neuron_ID)

Variables:

NormalizedNeuronalActivity: Mean Z-transformed PSTH (100-300 ms after target onset)

Scene: Fixed effect (levels: 1-4)

Value: Fixed effect (levels: good, bad)

Direction: Fixed effect (levels: contralateral, ipsilateral)

monkey_ID, Neuron_ID: Random effects

Multiple comparisons:

6 pairwise comparisons (good vs. bad, contralateral vs. ipsilateral) Bonferroni-corrected threshold: α = 0.05/6

### 3. Linear mixed-effects model for neuronal activity at saccade onset (Figure S2C)

Model objective: To examine neuronal activity patterns aligned to saccade onset.

Model specifications:

Full Model: NormalizedNeuronalActivity ∼ Scene × Value × Direction + (1|monkey_ID) + (1|monkey_ID: Neuron_ID)

Null Model: NormalizedNeuronalActivity ∼ (1|monkey_ID) + (1|monkey_ID: Neuron_ID)

Variables:

NormalizedNeuronalActivity: Mean Z-transformed PSTH (from 150 ms before to 50 ms after saccade onset)

Scene: Fixed effect (levels: 1-4)

Value: Fixed effect (levels: good, bad)

Direction: Fixed effect (levels: contralateral, ipsilateral)

monkey_ID, Neuron_ID: Random effects

Multiple comparisons:

6 pairwise comparisons (good vs. bad, contralateral vs. ipsilateral) Bonferroni-corrected threshold: α = 0.05/6

### 4. Bayesian linear mixed-effects model for neural activity and reaction time (Figure S2D)

Model objective: To examine the relationship between neural activity and saccadic reaction times.

Model specifications:

NormalizedRT ∼ NormalizedNeuronalActivity + (1|monkey_ID) + (1|monkey_ID: Neuron_ID)

Variables:

NormalizedRT: Standardized reaction times (mean = 0, SD = 1)

NormalizedNeuronalActivity: Mean Z-transformed PSTH (from 150 ms before to 50 ms after saccade onset)

monkey_ID, Neuron_ID: Random effects

MCMC specifications:

Chains: 4

Iterations: 50,000 per chain

Warmup: 5,000 per chain

Convergence criteria: Rhat = 1.00

These parameters (number of chains, total iterations, and warmup period) were selected to ensure adequate, effective sample sizes, stable parameter estimates, and reliable convergence diagnostics, thereby enhancing the credibility of our posterior inferences.

Initial model fitting using mean RT values failed to converge. As a result, RTs were standardized (mean = 0, SD = 1) prior to model estimation.

Model convergence was evaluated using the potential scale reduction factor (Rhat) and effective sample sizes (ESS). All Rhat values were 1.00, indicating successful convergence. Posterior distributions were summarized by their mean, 95% credible intervals (CIs), and probabilities that the regression coefficient for NormalizedFR (β_1_) was greater than 0. Statistical significance was inferred if the 95% CI did not include 0 and the probability of a positive coefficient exceeded 95%. All analyses were performed separately for four conditions: contralateral good objects, contralateral bad objects, ipsilateral good objects, and ipsilateral bad objects.

### 5. Linear mixed-effects model for neuronal activity during choice rejection and fixation (Figures 3E and H)

Model objective: To compare neuronal activity patterns between different rejection strategies and across choice and fixation tasks.

Model specifications:

Full Model: NormalizedNeuronalActivity ∼ Condition × Direction + (1|monkey_ID) + (1|monkey_ID: Neuron_ID)

Null Model: NormalizedNeuronalActivity ∼ (1|monkey_ID) + (1|monkey_ID: Neuron_ID)

Variables:

NormalizedNeuronalActivity: Mean Z-transformed PSTH (100-300 ms post-target)

Condition: Fixed effect (levels: return (choice task), stay (choice task), good (fixation task), bad (fixation task))

Direction: Fixed effect (levels: contralateral, ipsilateral)

Multiple comparisons:

6 pairwise comparisons between conditions

Bonferroni-corrected threshold: α = 0.05/6

### 6. Generalized linear mixed-effects model for saccade reaction times after injection (Figure 4A)

Model objective: To investigate the effects of glutamatergic antagonist injection on saccadic reaction times.

Model specifications:

Full Model: MedianSaccadeReactionTimes ∼ Injection × PrePost × Value × Direction + (1|monkey_ID) + (1|monkey_ID: Session_ID)

Null Model: MedianSaccadeReactionTimes ∼ (1|monkey_ID) + (1|monkey_ID: Session_ID)

Variables:

MedianSaccadeReactionTimes: Median reaction time per condition

Injection: Fixed effect (levels: antagonist, saline)

PrePost: Fixed effect (levels: preinjection, postinjection)

Value: Fixed effect (levels: good, bad)

Direction: Fixed effect (levels: contralateral, ipsilateral)

Session_ID: Random effect identifying individual injection sessions

Distribution: Poisson

Rationale for distribution choice: Reaction times are non-negative count data characterized by a right-skewed distribution.

Multiple comparisons:

8 pairwise comparisons (pre vs. post for each condition)

Bonferroni-corrected threshold: α = 0.05/8

### 7. Generalized linear mixed-effects model for chosen action rate after injection (Figure 4B)

Model objective: To investigate the impact of glutamatergic antagonist injection on action selection for bad objects.

Model specifications:

Full Model: ChosenActionRate ∼ Injection × PrePost × Value × Direction + (1|monkey_ID) + (1|monkey_ID: Session_ID), weights = (total trial count)

Null Model: ChosenActionRate ∼ (1|monkey_ID) + (1|monkey_ID: Session_ID), weights = (total trial count)

Variables:

ChosenActionRate: Proportion of selected actions

Injection, PrePost, Value, Direction: Fixed effects as described above total_trial_count: Weights to account for different numbers of trials

Distribution: Binomial

Rationale for distribution choice: Analysis of proportional data with binary outcomes

Multiple comparisons:

8 pairwise comparisons

Bonferroni-corrected threshold: α = 0.05/8

### 8. Generalized linear mixed-effects model for fixation break error rate after injection (Figure 4C)

Model objective: To examine how glutamatergic antagonist injection affects the ability to suppress reflexive saccades.

Model specifications:

Full Model: FixBreakErrorRate ∼ Injection × PrePost × Value × Direction + (1|monkey_ID) + (1|monkey_ID: Session_ID), weights = (total trial count)

Null Model: FixBreakErrorRate ∼ (1|monkey_ID) + (1|monkey_ID: Session_ID), weights = (total trial count)

Variables:

FixBreakErrorRate: Proportion of fixation break errors All other variables, as defined above

Distribution: Binomial

Rationale for distribution choice: Analysis of error rate data with binary outcomes.

Multiple comparisons:

8 pairwise comparisons

Bonferroni-corrected threshold: α = 0.05/8

## Acknowledgments

This research was supported by the Intramural Research Program at the National Institutes of Health, National Eye Institute (1ZIA EY000415). MRI scanning was conducted in the Neurophysiology Imaging Facility Core (National Institute of Mental Health, National Institute of Neurological Disorders and Stroke, and National Eye Institute). We thank Richard Krauzlis, the lab chief of the laboratory of Sensorimotor Research at the National Eye Institute for his support as a co-mentor. We also thank D. Parker, H. Warnock, G. Tansey, K. Allen-Worthington, A.M. Nichols, D. Yochelson, J. Fuller-Deets, and M. Robinson for technical assistance.

## Figures and Tables

**Fig. S1.**
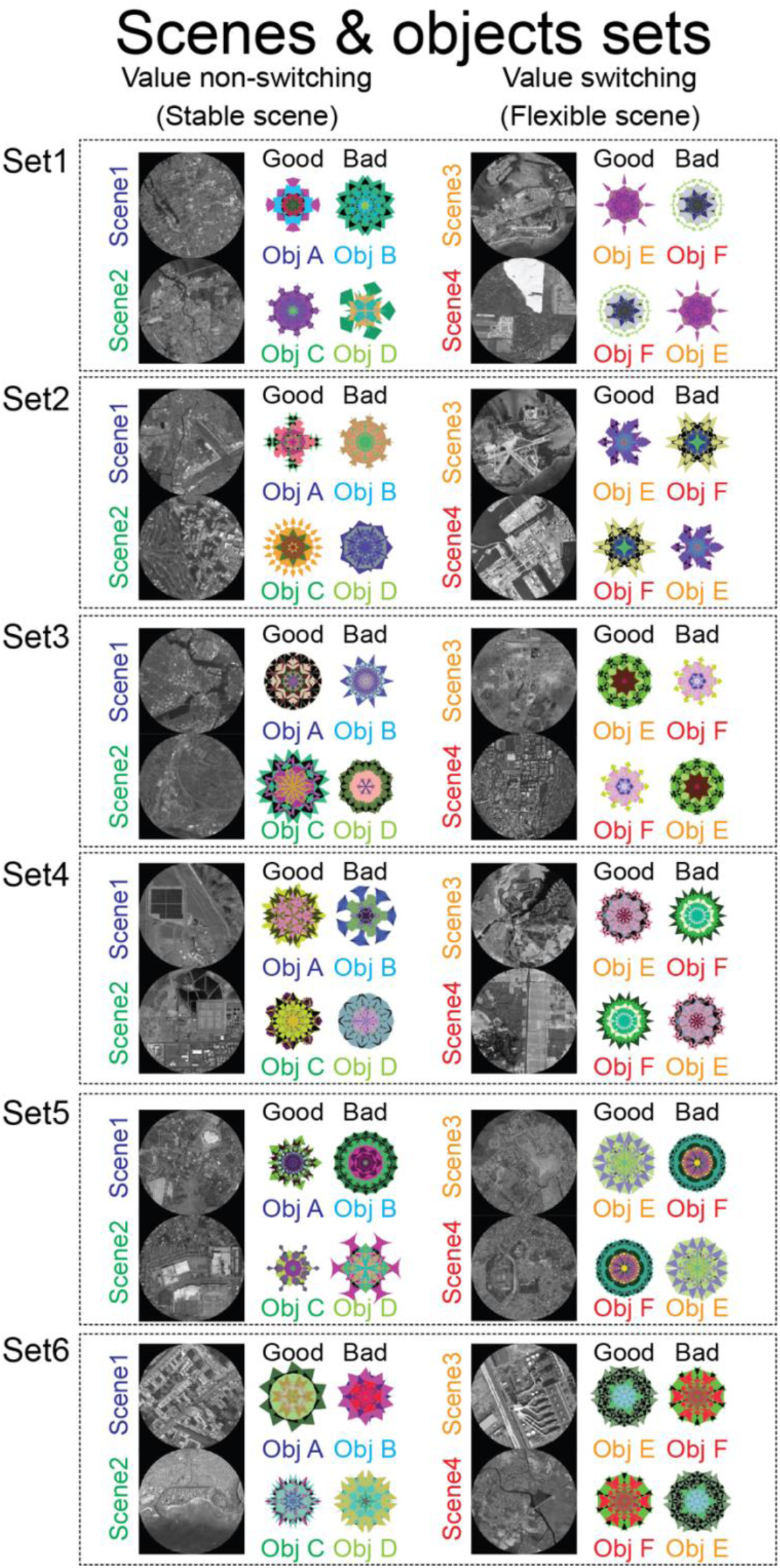
All sets of scenes 1-4 and good and bad objects for the choice task.

**Fig. S2.**
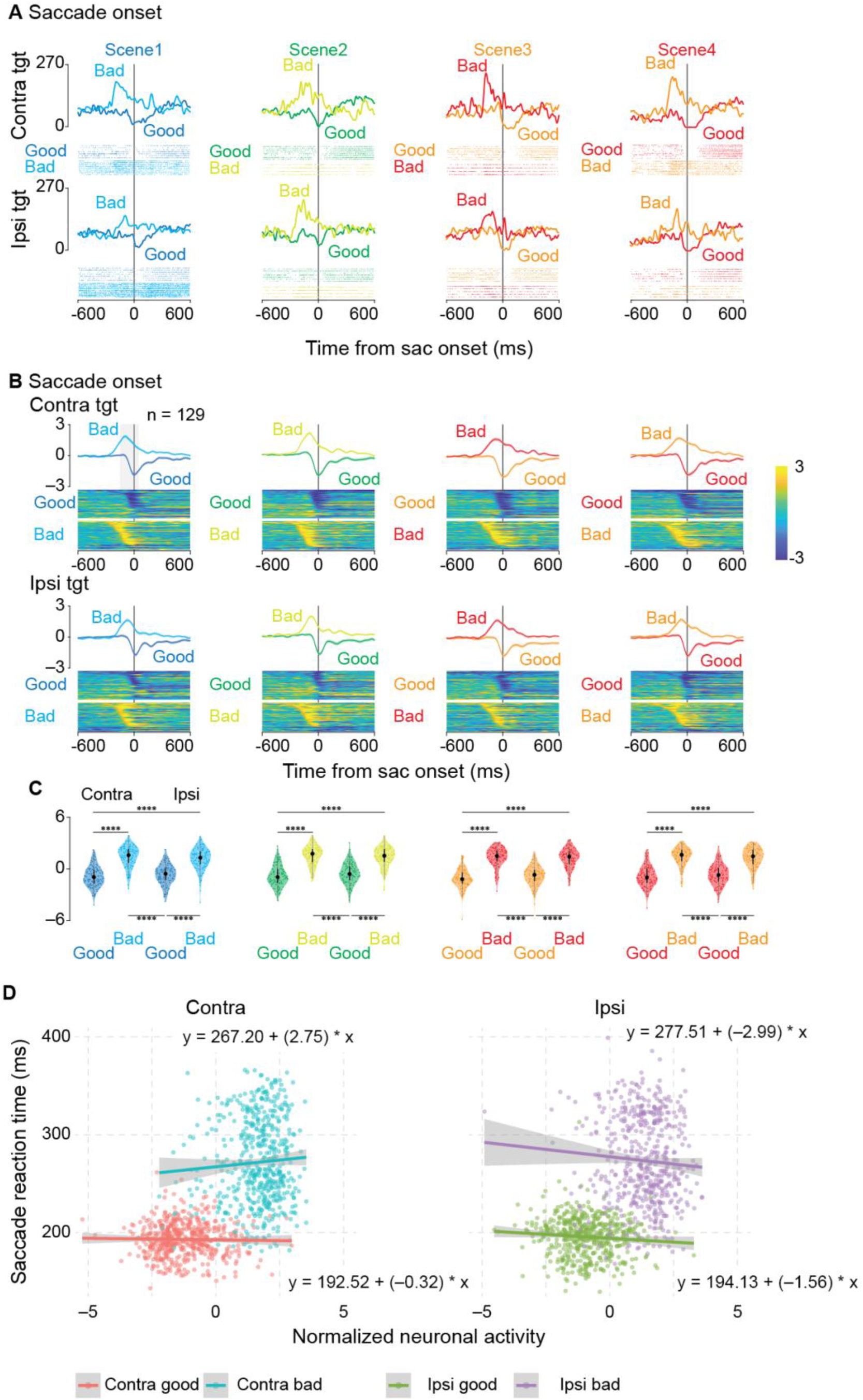
Neuronal activity aligned to saccade onset during the choice task (A) Activity of the same representative SNr neuron shown in Figure 2B, now aligned to saccade onset. Data are displayed separately for contralateral (upper) and ipsilateral (lower) target presentations, with different colors representing responses to good and bad objects in each scene. (B) Population activity aligned to saccade onset (n = 129 neurons). Upper panels display mean firing rates for contralateral target presentations, while lower panels show those for ipsilateral presentations. Population averages (upper traces) and normalized single-neuron responses (heat maps) are shown for good and bad objects across all scenes. (C) Distribution of neuronal responses. Violin plots quantify activity during a 200-ms window spanning from 150 ms before to 50 ms after saccade onset. Asterisks denote significant differences between conditions (*p < 0.05, **p < 0.01, ***p < 0.001, ****p < 0.0001, post-hoc pairwise t-tests with Bonferroni correction). (D) Relationship between normalized neuronal activity and saccadic reaction times for different target conditions. Scatter plots show individual trials with regression lines (colored lines) and 95% confidence intervals (gray shading). Separate analyses are presented for contralateral good (red), contralateral bad (cyan), ipsilateral good (green), and ipsilateral bad (purple) targets. Regression equations are provided for each condition.

**Table S1.**
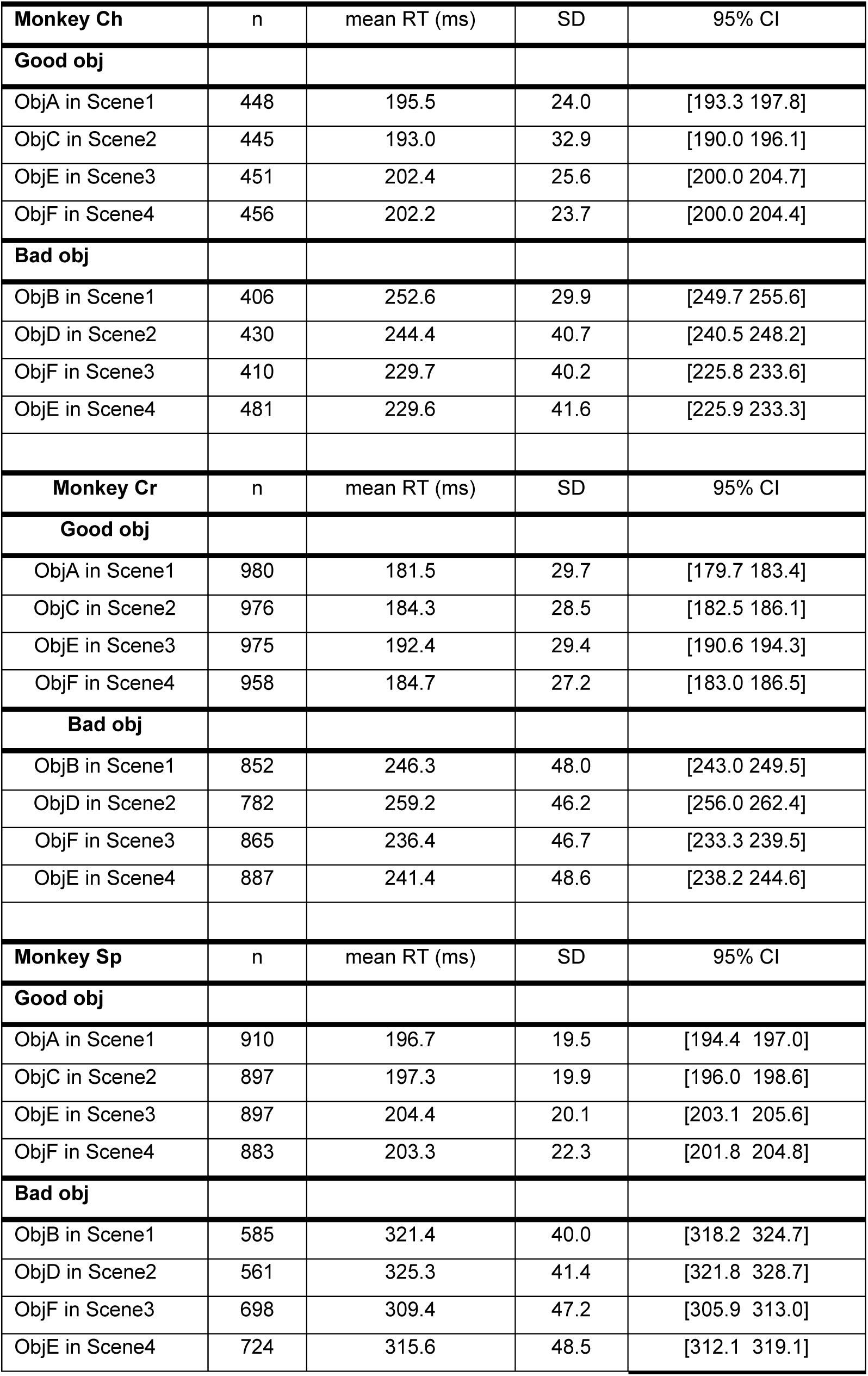
Saccade reaction times in each condition of each monkey.

| **Monkey Ch** | **n** | **mean RT (ms)** | **SD** | **95% CI** |
| --- | --- | --- | --- | --- |
| **Good obj** |  |  |  |  |
| ObjA in Scene1 | 448 | 195.5 | 24.0 | [193.3 197.8] |
| ObjC in Scene2 | 445 | 193.0 | 32.9 | [190.0 196.1] |
| ObjE in Scene3 | 451 | 202.4 | 25.6 | [200.0 204.7] |
| ObjF in Scene4 | 456 | 202.2 | 23.7 | [200.0 204.4] |
| **Bad obj** |  |  |  |  |
| ObjB in Scene1 | 406 | 252.6 | 29.9 | [249.7 255.6] |
| ObjD in Scene2 | 430 | 244.4 | 40.7 | [240.5 248.2] |
| ObjF in Scene3 | 410 | 229.7 | 40.2 | [225.8 233.6] |
| ObjE in Scene4 | 481 | 229.6 | 41.6 | [225.9 233.3] |
| **Monkey Cr** | **n** | **mean RT (ms)** | **SD** | **95% CI** |
| **Good obj** |  |  |  |  |
| ObjA in Scene1 | 980 | 181.5 | 29.7 | [179.7 183.4] |
| ObjC in Scene2 | 976 | 184.3 | 28.5 | [182.5 186.1] |
| ObjE in Scene3 | 975 | 192.4 | 29.4 | [190.6 194.3] |
| ObjF in Scene4 | 958 | 184.7 | 27.2 | [183.0 186.5] |
| **Bad obj** |  |  |  |  |
| ObjB in Scene1 | 852 | 246.3 | 48.0 | [243.0 249.5] |
| ObjD in Scene2 | 782 | 259.2 | 46.2 | [256.0 262.4] |
| ObjF in Scene3 | 865 | 236.4 | 46.7 | [233.3 239.5] |
| ObjE in Scene4 | 887 | 241.4 | 48.6 | [238.2 244.6] |
| **Monkey Sp** | **n** | **mean RT (ms)** | **SD** | **95% CI** |
| **Good obj** |  |  |  |  |
| ObjA in Scene1 | 910 | 196.7 | 19.5 | [194.4 197.0] |
| ObjC in Scene2 | 897 | 197.3 | 19.9 | [196.0 198.6] |
| ObjE in Scene3 | 897 | 204.4 | 20.1 | [203.1 205.6] |
| ObjF in Scene4 | 883 | 203.3 | 22.3 | [201.8 204.8] |
| **Bad obj** |  |  |  |  |
| ObjB in Scene1 | 585 | 321.4 | 40.0 | [318.2 324.7] |
| ObjD in Scene2 | 561 | 325.3 | 41.4 | [321.8 328.7] |
| ObjF in Scene3 | 698 | 309.4 | 47.2 | [305.9 313.0] |
| ObjE in Scene4 | 724 | 315.6 | 48.5 | [312.1 319.1] |

**Table S2.**
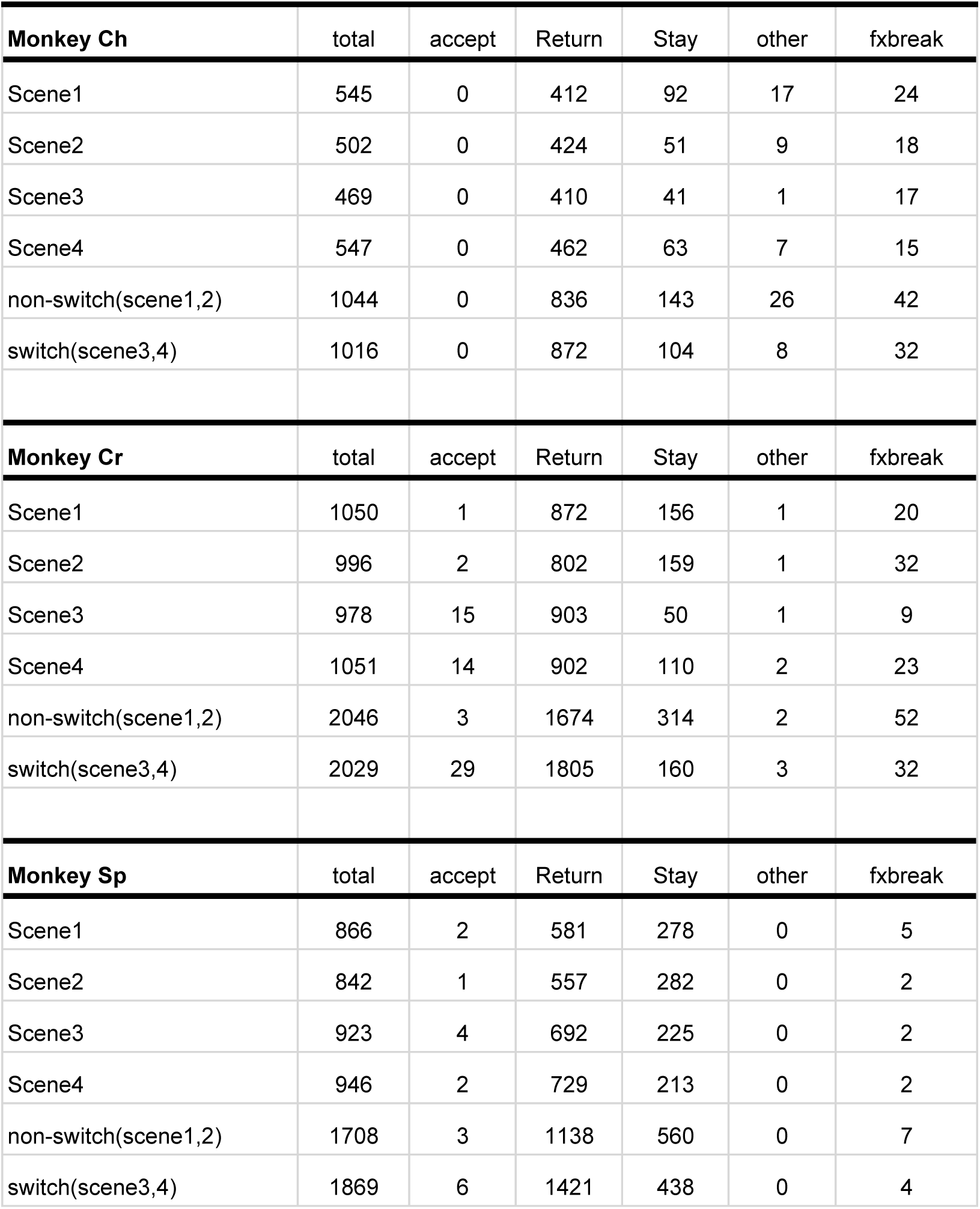
Counts of chosen actions for Bad objects.

**Table S3.**
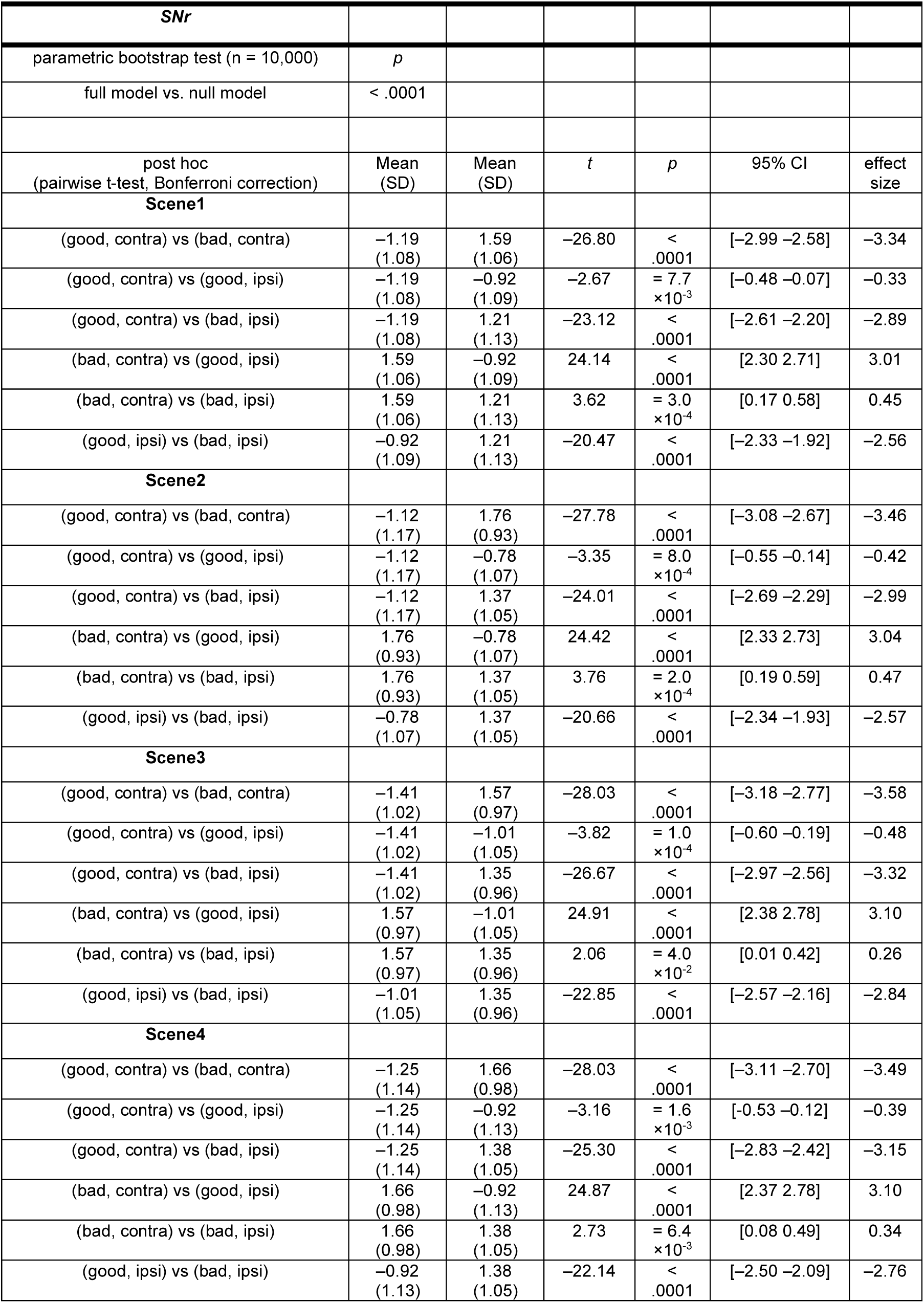
Summary of statistical test to compare the normalized neuronal activity of SNr neurons at target onset among conditions during choice task in Figure 2.

| **SNr** |  |  |  |  |  |  |
| --- | --- | --- | --- | --- | --- | --- |
| parametric bootstrap test (n = 10,000) | *p* |  |  |  |  |  |
| full model vs. null model | < .0001 |  |  |  |  |  |
| post hoc<br>(pairwise t-test, Bonferroni correction) | Mean<br>(SD) | Mean<br>(SD) | *t* | *p* | 95% CI | effect<br>size |
| **Scene1** |  |  |  |  |  |  |
| (good, contra) vs (bad, contra) | -1.19<br>(1.08) | 1.59<br>(1.06) | -26.80 | <<br>.0001 | [-2.99 -2.58] | -3.34 |
| (good, contra) vs (good, ipsi) | -1.19<br>(1.08) | -0.92<br>(1.09) | -2.67 | = 7.7<br>$\times 10^{-3}$ | [-0.48 -0.07] | -0.33 |
| (good, contra) vs (bad, ipsi) | -1.19<br>(1.08) | 1.21<br>(1.13) | -23.12 | <<br>.0001 | [-2.61 -2.20] | -2.89 |
| (bad, contra) vs (good, ipsi) | 1.59<br>(1.06) | -0.92<br>(1.09) | 24.14 | <<br>.0001 | [2.30 2.71] | 3.01 |
| (bad, contra) vs (bad, ipsi) | 1.59<br>(1.06) | 1.21<br>(1.13) | 3.62 | = 3.0<br>$\times 10^{-4}$ | [0.17 0.58] | 0.45 |
| (good, ipsi) vs (bad, ipsi) | -0.92<br>(1.09) | 1.21<br>(1.13) | -20.47 | <<br>.0001 | [-2.33 -1.92] | -2.56 |
| **Scene2** |  |  |  |  |  |  |
| (good, contra) vs (bad, contra) | -1.12<br>(1.17) | 1.76<br>(0.93) | -27.78 | <<br>.0001 | [-3.08 -2.67] | -3.46 |
| (good, contra) vs (good, ipsi) | -1.12<br>(1.17) | -0.78<br>(1.07) | -3.35 | = 8.0<br>$\times 10^{-4}$ | [-0.55 -0.14] | -0.42 |
| (good, contra) vs (bad, ipsi) | -1.12<br>(1.17) | 1.37<br>(1.05) | -24.01 | <<br>.0001 | [-2.69 -2.29] | -2.99 |
| (bad, contra) vs (good, ipsi) | 1.76<br>(0.93) | -0.78<br>(1.07) | 24.42 | <<br>.0001 | [2.33 2.73] | 3.04 |
| (bad, contra) vs (bad, ipsi) | 1.76<br>(0.93) | 1.37<br>(1.05) | 3.76 | = 2.0<br>$\times 10^{-4}$ | [0.19 0.59] | 0.47 |
| (good, ipsi) vs (bad, ipsi) | -0.78<br>(1.07) | 1.37<br>(1.05) | -20.66 | <<br>.0001 | [-2.34 -1.93] | -2.57 |
| **Scene3** |  |  |  |  |  |  |
| (good, contra) vs (bad, contra) | -1.41<br>(1.02) | 1.57<br>(0.97) | -28.03 | <<br>.0001 | [-3.18 -2.77] | -3.58 |
| (good, contra) vs (good, ipsi) | -1.41<br>(1.02) | -1.01<br>(1.05) | -3.82 | = 1.0<br>$\times 10^{-4}$ | [-0.60 -0.19] | -0.48 |
| (good, contra) vs (bad, ipsi) | -1.41<br>(1.02) | 1.35<br>(0.96) | -26.67 | <<br>.0001 | [-2.97 -2.56] | -3.32 |
| (bad, contra) vs (good, ipsi) | 1.57<br>(0.97) | -1.01<br>(1.05) | 24.91 | <<br>.0001 | [2.38 2.78] | 3.10 |
| (bad, contra) vs (bad, ipsi) | 1.57<br>(0.97) | 1.35<br>(0.96) | 2.06 | = 4.0<br>$\times 10^{-2}$ | [0.01 0.42] | 0.26 |
| (good, ipsi) vs (bad, ipsi) | -1.01<br>(1.05) | 1.35<br>(0.96) | -22.85 | <<br>.0001 | [-2.57 -2.16] | -2.84 |
| **Scene4** |  |  |  |  |  |  |
| (good, contra) vs (bad, contra) | -1.25<br>(1.14) | 1.66<br>(0.98) | -28.03 | <<br>.0001 | [-3.11 -2.70] | -3.49 |
| (good, contra) vs (good, ipsi) | -1.25<br>(1.14) | -0.92<br>(1.13) | -3.16 | = 1.6<br>$\times 10^{-3}$ | [-0.53 -0.12] | -0.39 |
| (good, contra) vs (bad, ipsi) | -1.25<br>(1.14) | 1.38<br>(1.05) | -25.30 | <<br>.0001 | [-2.83 -2.42] | -3.15 |
| (bad, contra) vs (good, ipsi) | 1.66<br>(0.98) | -0.92<br>(1.13) | 24.87 | <<br>.0001 | [2.37 2.78] | 3.10 |
| (bad, contra) vs (bad, ipsi) | 1.66<br>(0.98) | 1.38<br>(1.05) | 2.73 | = 6.4<br>$\times 10^{-3}$ | [0.08 0.49] | 0.34 |
| (good, ipsi) vs (bad, ipsi) | -0.92<br>(1.13) | 1.38<br>(1.05) | -22.14 | <<br>.0001 | [-2.50 -2.09] | -2.76 |

**Table S4.** Summary of statistical test to compare the normalized neuronal activity of SNr neurons at saccade onset among conditions during choice task in Figure S2.

| **SNr** |  |  |  |  |  |  |
| --- | --- | --- | --- | --- | --- | --- |
| parametric bootstrap test (n = 10,000) | *p* |  |  |  |  |  |
| full model vs. null model | < .0001 |  |  |  |  |  |
| post hoc<br>(pairwise t-test, Bonferroni correction) | Mean<br>(SD) | Mean<br>(SD) | *t* | *p* | 95% CI | effect<br>size |
| **Scene1** |  |  |  |  |  |  |
| (good, contra) vs (bad, contra) | -0.87<br>(1.17) | 1.39<br>(1.21) | -20.16 | <<br>.0001 | [-2.48 -2.04] | -2.52 |
| (good, contra) vs (good, ipsi) | -0.87<br>(1.17) | -0.65<br>(1.71) | -1.97 | = 4.9<br>$\times 10^{-2}$ | [-0.44 -0.01] | -0.25 |
| (good, contra) vs (bad, ipsi) | -0.87<br>(1.17) | 1.17<br>(1.25) | -18.16 | <<br>.0001 | [-2.26 -1.82] | -2.27 |
| (bad, contra) vs (good, ipsi) | 1.39<br>(1.21) | -0.65<br>(1.71) | 18.19 | <<br>.0001 | [1.82 2.26] | 2.27 |
| (bad, contra) vs (bad, ipsi) | 1.39<br>(1.21) | 1.17<br>(1.25) | 1.96 | = 5.1<br>$\times 10^{-2}$ | [-0.01 0.44] | 0.25 |
| (good, ipsi) vs (bad, ipsi) | -0.65<br>(1.71) | 1.17<br>(1.25) | -16.20 | <<br>.0001 | [-2.04 -1.60] | -2.03 |
| **Scene2** |  |  |  |  |  |  |
| (good, contra) vs (bad, contra) | -0.80<br>(1.21) | 1.54<br>(1.11) | -20.93 | <<br>.0001 | [-2.56 -2.13] | -2.61 |
| (good, contra) vs (good, ipsi) | -0.80<br>(1.21) | -0.54<br>(1.18) | -2.36 | = 1.8<br>$\times 10^{-2}$ | [-0.48 -0.04] | -0.29 |
| (good, contra) vs (bad, ipsi) | -0.80<br>(1.21) | 1.39<br>(1.18) | -19.51 | <<br>.0001 | [-2.41 -1.97] | -2.43 |
| (bad, contra) vs (good, ipsi) | 1.54<br>(1.11) | -0.70<br>(1.13) | 18.57 | <<br>.0001 | [1.86 2.30] | 2.31 |
| (bad, contra) vs (bad, ipsi) | 1.54<br>(1.11) | 1.39<br>(1.18) | 1.38 | = .167 | [-0.07 0.38] | 0.17 |
| (good, ipsi) vs (bad, ipsi) | -0.70<br>(1.13) | 1.39<br>(1.18) | -17.15 | <<br>.0001 | [-2.15 -1.71] | -2.14 |
| **Scene3** |  |  |  |  |  |  |
| (good, contra) vs (bad, contra) | -1.11<br>(1.10) | 1.35<br>(1.10) | -21.93 | <<br>.0001 | [-2.68 -2.24] | -2.73 |
| (good, contra) vs (good, ipsi) | -1.11<br>(1.10) | -0.80<br>(1.21) | -2.77 | = 6.0<br>$\times 10^{-3}$ | [-0.53 -0.09] | -0.35 |
| (good, contra) vs (bad, ipsi) | -1.11<br>(1.10) | 1.26<br>(1.09) | -21.11 | <<br>.0001 | [-2.59 -2.15] | -2.63 |
| (bad, contra) vs (good, ipsi) | 1.35<br>(1.10) | -0.80<br>(1.21) | 19.16 | <<br>.0001 | [1.93 2.37] | 2.39 |
| (bad, contra) vs (bad, ipsi) | 1.35<br>(1.10) | 1.26<br>(1.09) | 0.82 | = .410 | [-0.13 0.31] | 0.10 |
| (good, ipsi) vs (bad, ipsi) | -0.80<br>(1.21) | 1.26<br>(1.09) | -18.34 | <<br>.0001 | [-2.27 -1.83] | -2.28 |
| **Scene4** |  |  |  |  |  |  |
| (good, contra) vs (bad, contra) | -0.87<br>(1.18) | 1.43<br>(1.07) | -20.47 | <<br>.0001 | [-2.51 -2.07] | -2.55 |
| (good, contra) vs (good, ipsi) | -0.87<br>(1.18) | -0.68<br>(1.18) | -1.70 | = 9.0<br>$\times 10^{-2}$ | [-0.41 -0.03] | -0.21 |
| (good, contra) vs (bad, ipsi) | -0.87<br>(1.18) | 1.30<br>(1.16) | -19.31 | <<br>.0001 | [-2.38 -1.94] | -2.40 |
| (bad, contra) vs (good, ipsi) | 1.43<br>(1.07) | -0.68<br>(1.18) | 18.78 | <<br>.0001 | [1.88 2.32] | 2.34 |
| (bad, contra) vs (bad, ipsi) | 1.43<br>(1.07) | 1.30<br>(1.16) | 1.17 | = .244 | [-0.09 0.35] | 0.15 |
| (good, ipsi) vs (bad, ipsi) | -0.68<br>(1.18) | 1.30<br>(1.16) | -17.61 | <<br>.0001 | [-2.19 -1.75] | -2.19 |

**Table S5.** Summary of statistical test to examine the relationship between the normalized neuronal activity and normalized saccade reaction times during choice task in Figure S2.

| Condition | Mean ($\beta$)<br>for<br>NormalizedFR | 95%<br>Credible<br>Interval<br>for $\beta$ | Probability<br>$\beta > 0$ | SD<br>(Monkey) | SD<br>(Neuron) | Residual<br>SD | Rhat<br>($\beta$) | Effective<br>Sample<br>Size ($\beta$) |
| --- | --- | --- | --- | --- | --- | --- | --- | --- |
| Contralateral<br>Good | 0.018 | [-0.05,<br>0.08] | 0.70 | 0.77 | 0.58 | 0.79 | 1.00 | 225328 |
| Contralateral<br>Bad | 0.075 | [0.01,<br>0.14] | 0.99 | 0.27 | 0.82 | 0.59 | 1.00 | 197998 |
| Ipsilateral<br>Good | -0.043 | [-0.11,<br>0.02] | 0.11 | 0.54 | 0.67 | 0.73 | 1.00 | 202677 |
| Ipsilateral<br>Bad | 0.010 | [-0.05,<br>0.06] | 0.57 | 0.33 | 0.84 | 0.56 | 1.00 | 144892 |

**Table S6.** Summary of statistical test to compare the normalized neuronal activity of SNr neurons among Return, Stay during choice task, and fixation task in Figure 3.

| *SNr* |  |  |  |  |  |  |
| --- | --- | --- | --- | --- | --- | --- |
| parametric bootstrap test (n = 10,000) | *p* |  |  |  |  |  |
| full model vs. null model | < .001 |  |  |  |  |  |
| post hoc<br>(pairwise t-test, Bonferroni correction) | Mean (SD) | Mean (SD) | *t* | *p* | 95% CI | effect size |
| Contra |  |  |  |  |  |  |
| (Return, choice) vs (Stay, choice) | 1.59 (1.06) | 1.40 (1.02) | 1.83 | = $6.7 \times 10^{-2}$ | [-0.01 0.42] | 0.25 |
| (Return, choice) vs (Good, fixation) | 1.59 (1.06) | 1.31 (1.06) | -2.71 | = $6.8 \times 10^{-3}$ | [-0.51 -0.08] | -0.37 |
| (Return, choice) vs (Bad, fixation) | 1.59 (1.06) | 1.57 (0.84) | -0.29 | = $7.7 \times 10^{-1}$ | [-0.24 0.18] | -0.04 |
| (Stay, choice) vs (Good, fixation) | 1.40 (1.02) | 1.31 (1.06) | -0.78 | = $4.3 \times 10^{-1}$ | [-0.32 0.14] | -0.11 |
| (Stay, choice) vs (Bad, fixation) | 1.40 (1.02) | 1.57 (0.84) | 1.44 | = $1.5 \times 10^{-1}$ | [-0.06 0.40] | 0.21 |
| (Good, fixation) vs (Bad, fixation) | 1.31 (1.06) | 1.57 (0.84) | 2.30 | = $2.1 \times 10^{-2}$ | [0.04 0.48] | 0.33 |
| Ipsi |  |  |  |  |  |  |
| (Return, choice) vs (Stay, choice) | 1.21 (1.13) | 1.17 (0.87) | 0.76 | = $7.6 \times 10^{-1}$ | [-0.18 0.25] | 0.04 |
| (Return, choice) vs (Good, fixation) | 1.21 (1.13) | 0.90 (0.82) | -3.05 | = $2.3 \times 10^{-3}$ | [-0.54 -0.12] | -0.42 |
| (Return, choice) vs (Bad, fixation) | 1.21 (1.13) | 0.51 (0.85) | -6.65 | < .0001 | [-0.93 -0.51] | -0.90 |
| (Stay, choice) vs (Good, fixation) | 1.17 (0.87) | 0.90 (0.82) | -2.51 | = $1.2 \times 10^{-2}$ | [-0.53 -0.06] | -0.37 |
| (Stay, choice) vs (Bad, fixation) | 1.17 (0.87) | 0.51 (0.85) | -5.80 | < .0001 | [-0.92 -0.45] | -0.86 |
| (Good, fixation) vs (Bad, fixation) | 0.90 (0.82) | 0.51 (0.85) | -3.44 | = $6.0 \times 10^{-4}$ | [-0.61 -0.17] | -0.49 |

**Table S7.** Summary of statistical test to compare the effects of CPP + NBQX injection into SNr during choice task in Figure 4.

| *Injection during choice task* |  |  |  |  |
| --- | --- | --- | --- | --- |
| parametric bootstrap test (n = 10,000) | *p* |  |  |  |
| full model vs. null model | < .0001 |  |  |  |
| post hoc<br>(pairwise t-test, Bonferroni correction) | Mean (ms)<br>(SD) | Mean (ms)<br>(SD) | *z* | *p* |
| CPP+NBQX Contra Good pre vs. post | 196.25<br>(9.11) | 171.42<br>(12.17) | -4.48 | < .0001 |
| Saline Contra Good pre vs. post | 194.17<br>(8.68) | 191.08<br>(7.81) | -0.54 | = .58 |
| CPP+NBQX Contra Bad pre vs. post | 267.75<br>(18.52) | 181.42<br>(17.83) | -14.01 | < .0001 |
| Saline Contra Bad pre vs. post | 271.42<br>(17.73) | 264.42<br>(22.38) | -1.05 | = .29 |
| CPP+NBQX Ipsi Good pre vs. post | 199.25<br>(10.17) | 214.92<br>(22.70) | 2.67 | = 7.7<br>×10<sup>-3</sup> |
| Saline Ipsi Good pre vs. post | 199.83<br>(8.83) | 195.92<br>(8.27) | -0.68 | = .50 |
| CPP+NBQX Ipsi Bad pre vs. post | 282.58<br>(31.80) | 280.08<br>(58.68) | -0.37 | = .72 |
| Saline Ipsi Bad pre vs. post | 288.75<br>(36.97) | 274.83<br>(38.89) | -2.03 | = 4.2<br>×10<sup>-2</sup> |

**Table S8.** Summary of statistical test to compare the effects of CPP + NBQX injection into SNr while monkeys chose actions for Bad object during Choice task in Figure 4.

| **Injection into SNr during Choice task** |  |  |  |  |
| --- | --- | --- | --- | --- |
| Accept Bad object |  |  |  |  |
| parametric bootstrap test (n = 10,000) | *p* |  |  |  |
| full model vs. null model | = 1.00 |  |  |  |
| Return for Bad object |  |  |  |  |
| parametric bootstrap test (n = 10,000) | *p* |  |  |  |
| full model vs. null model | < .001 |  |  |  |
| post hoc<br>(pairwise t-test, Bonferroni correction) | Mean (%)<br>(SD) | Mean (%)<br>(SD) | *z* | *p* |
| CPP+NBQX<br>Return Contra Bad pre vs. post | 80.94<br>(15.30) | 88.73<br>(10.99) | 3.41 | = $7.0 \times 10^{-4}$ |
| Saline<br>Return Contra Good pre vs. post | 78.93<br>(13.43) | 77.99<br>(14.15) | -0.63 | = .53 |
| CPP+NBQX<br>Return Ipsi Bad pre vs. post | 72.04<br>(20.47) | 48.38<br>(24.41) | -8.14 | < .0001 |
| Saline<br>Return Ipsi Bad pre vs. post | 68.13<br>(17.52) | 70.48<br>(19.81) | 0.28 | = .78 |
| Stay for Bad object |  |  |  |  |
| parametric bootstrap test (n = 10,000) | *p* |  |  |  |
| full model vs. null model | < .001 |  |  |  |
| post hoc<br>(pairwise t-test, Bonferroni correction) | Mean (%)<br>(SD) | Mean (%)<br>(S.D.) | *z* | *p* |
| CPP+NBQX<br>Return Contra Bad pre vs. post | 19.51<br>(15.31) | 3.70<br>(8.61) | -6.47 | < .0001 |
| Saline<br>Return Contra Good pre vs. post | 21.07<br>(13.42) | 21.68<br>(13.82) | 0.38 | = .70 |
| CPP+NBQX<br>Return Ipsi Bad pre vs. post | 27.78<br>(20.47) | 50.46<br>(23.36) | 7.90 | < .0001 |
| Saline<br>Return Ipsi Bad pre vs. post | 31.79<br>(17.55) | 29.37<br>(19.58) | -0.40 | = .69 |

**Table S9.** Summary of statistical test to compare the effects of CPP + NBQX injection into *SNr during fixation task in Figure 4*.

| *Injection into SNr during fixation task* |  |  |  |  |
| --- | --- | --- | --- | --- |
| parametric bootstrap test (n = 10,000) | *p* |  |  |  |
| full model vs. null model | < .05 |  |  |  |
| post hoc<br>(pairwise t-test, Bonferroni correction) | Mean (%)<br>(SD) | Mean (%)<br>(SD) | *t* | *z* |
| CPP+NBQX Contra Good pre vs. post | 2.62<br>(4.23) | 63.97<br>(25.43) | 14.85 | < .0001 |
| Saline Contra Good pre vs. post | 1.85<br>(2.12) | 2.57<br>(3.40) | 0.65 | = .51 |
| CPP+NBQX Contra Bad pre vs. post | 1.86<br>(2.47) | 52.51<br>(24.40) | 12.10 | < .0001 |
| Saline Contra Bad pre vs. post | 0.93<br>(1.89) | 0.75<br>(1.54) | -0.08 | = .94 |
| CPP+NBQX Ipsi Good pre vs. post | 2.43<br>(3.32) | 0.17<br>(0.58) | -2.18 | = .03 |
| Saline Ipsi Good pre vs. post | 0.51<br>(0.97) | 0.67<br>(1.21) | 0.21 | = .84 |
| CPP+NBQX Ipsi Bad pre vs. post | 1.22<br>(2.50) | 0.80<br>(1.19) | -0.79 | = .43 |
| Saline Ipsi Bad pre vs. post | 0.24<br>(0.82) | 0.22<br>(0.76) | 0.06 | = .95 |

